# Neural birth time and somatosensory circuit assembly are linked by Robo3 regulation of dendrite morphology

**DOI:** 10.64898/2026.05.29.728185

**Authors:** Jake E. Henderson, Nicolas Zerda Afanador, Emma Carlson, Ellie S. Heckscher

## Abstract

Neural circuit wiring requires a remarkable level of precision, as thousands of neurons generate millions of synapses in distinct configurations. While neural circuits are known to be shaped by both neural birth timing and multiple classes of guidance molecules, the relationship between the two and how they coordinate circuit assembly has yet to be evaluated. Leveraging the well-defined stem cell lineages of the Drosophila embryonic nerve cord, we investigate how Roundabout (Robo) guidance receptors are regulated by neural birth timing to establish neural morphology and function for circuit assembly. Our data reveal that Robo receptor expression in neurons is tightly linked to neural birth timing and controlled by temporal transcription factors. Notably, we identify a Robo3 guidance receptor code that is pivotal for assembly of somatosensory circuits, distinguishing stretch-detecting PNS neurons and late-born EL interneurons (lacking Robo3) from vibration detecting PNS neurons and early-born EL interneurons (expressing Robo3). These insights propose a shared guidance receptor code connecting neural birth timing to circuit assembly, providing a molecular foundation for developmental wiring specificity.

**Highlights:**

1. Robo3 is enriched in early-born neurons and requires Pdm and Castor for normal expression
2. Classic temporal identity markers and Robo3 are differentially regulated
3. In interneurons, Robo3 is required for dendrite positioning and branching
4. A shared Robo code is used by synaptic partners from different developmental origins

## Introduction

During neural circuit assembly, proper wiring requires thousands of neurons to form millions of synapses in distinct configurations, all guided by axon guidance molecules^1^. Axon guidance ligand-receptor systems have the hallmarks of a redundant molecular code: multiple ligands pattern a particular axis, and for each ligand, multiple receptors exist^1–8^. One example of this redundant molecular code is the patterning of the medio-lateral axis in Drosophila. At the midline, Netrin and Slit ligands are secreted^2,5,9–12^. Neurons respond to each ligand with different receptor families: Frazzled/Dcc and Unc-5 for Netrin, and Roundabouts 1, 2, 3 for Slit^2,10,11,13–16^. Homologous molecules pattern the mediolateral axis across phyla^17–19^. Redundancy makes the molecular code robust against mutations or changes in gene number^1,3,5,10,20,21^. However, redundancy introduces complexity, making it difficult to understand how axon guidance molecules are used in circuit assembly.

How axon guidance cues are used in circuit assembly is just beginning to be understood. For example, Slit2/Robo1 signaling was recently shown to refine activity of direction selective circuitry, thereby maintaining image stabilization during behavior^22^. But our understanding still falls short of knowing how axon guidance cues are used to assemble ensembles of neurons into specific networks of connectivity. To assemble any type of circuit, neurons from different locations must project to a given region in the developing CNS before they can form synapses^8,23,24^. The redundant ligand/receptor code means that any given neuron has a vast number of potential combinations it could use to get to a particular location. The extent to which synaptic partners share a code or take advantage of the redundancy is unknown.

Furthermore, in many systems, circuit assembly is associated with neuronal birth timing^25–43^. Ultimately, birth timing must regulate circuit wiring by impacting the expression and/or activity of axon guidance receptors/ligands. Expression of the Robo receptors in Drosophila, for example, can differ both in temporal dynamics and spatial pattern^16^. A handful of terminal selector transcription factors (e.g., Eve, Hb9, Islet) are known to regulate the expression of axon guidance molecules as part of terminal differentiation^17,44–49^. But what we know about transcriptional regulation of the redundant axon guidance code neither reveals nor explains any association between circuit assembly and birth timing.

In this study, to understand the role of Robo receptors in circuit assembly, we used a well-established example of timing in circuit wiring, the EL somatosensory processing circuits in the nerve cord of Drosophila larvae (Figure 1). In this model, one neural stem cell, Neuroblast 3-3 (NB3-3), gives rise to two sets of Even-skipped Lateral (EL) neurons—five early-born ELs, followed by six late-born ELs(Figure 1)^31,50^. Both types of ELs process somatosensory stimuli, but the modality differs by birth time: early-born ELs encode mechanical/vibrational stimuli, whereas late-born ELs encode proprioceptive/self-movement stimuli^31,51^. The underlying circuit architectures are known: early-born ELs synapse with vibration-sensitive chordotonal (CHO) sensory neurons, whereas late-born ELs synapse with stretch-sensitive dorsal bipolar dendrite (DBD) and other proprioceptive sensory neurons (Figure 1A)^31,41^. The axon guidance molecules that underlie the establishment of these circuits are partially understood: In mature circuits, CHO and proprioceptors like DBD have axons that terminate at different mediolateral positions within the CNS. During embryogenesis, CHO express both Robo1 and Robo3, whereas DBD express Robo1 but not Robo3 (Figure 1B)^52^. Thus, Robo3 expression is the key molecular switch that repels CHO axons farther from the midline. In contrast, the axon guidance cues that regulate the EL side of these circuits remain unknown. Early-born ELs must send their dendrites to a more lateral position in the CNS to meet CHO axons, while late-born ELs must send their dendrites to a more medial position to meet DBD axons. So, early-born ELs must experience different amounts of attraction and/or repulsion from the midline compared to the late-born ELs. One hypothesis is that Robo3 expression might also act as a key switch in ELs—with early-born ELs requiring Robo3 for proper dendrite placement and vibrational encoding. However, the extent to which Robo3 is differentially expressed in neurons of different birth times, including ELs of the NB3-3, is unknown (Figure 1C), as is the role of Robo3 in the EL somatosensory circuit structure and function.

**Figure 1.**
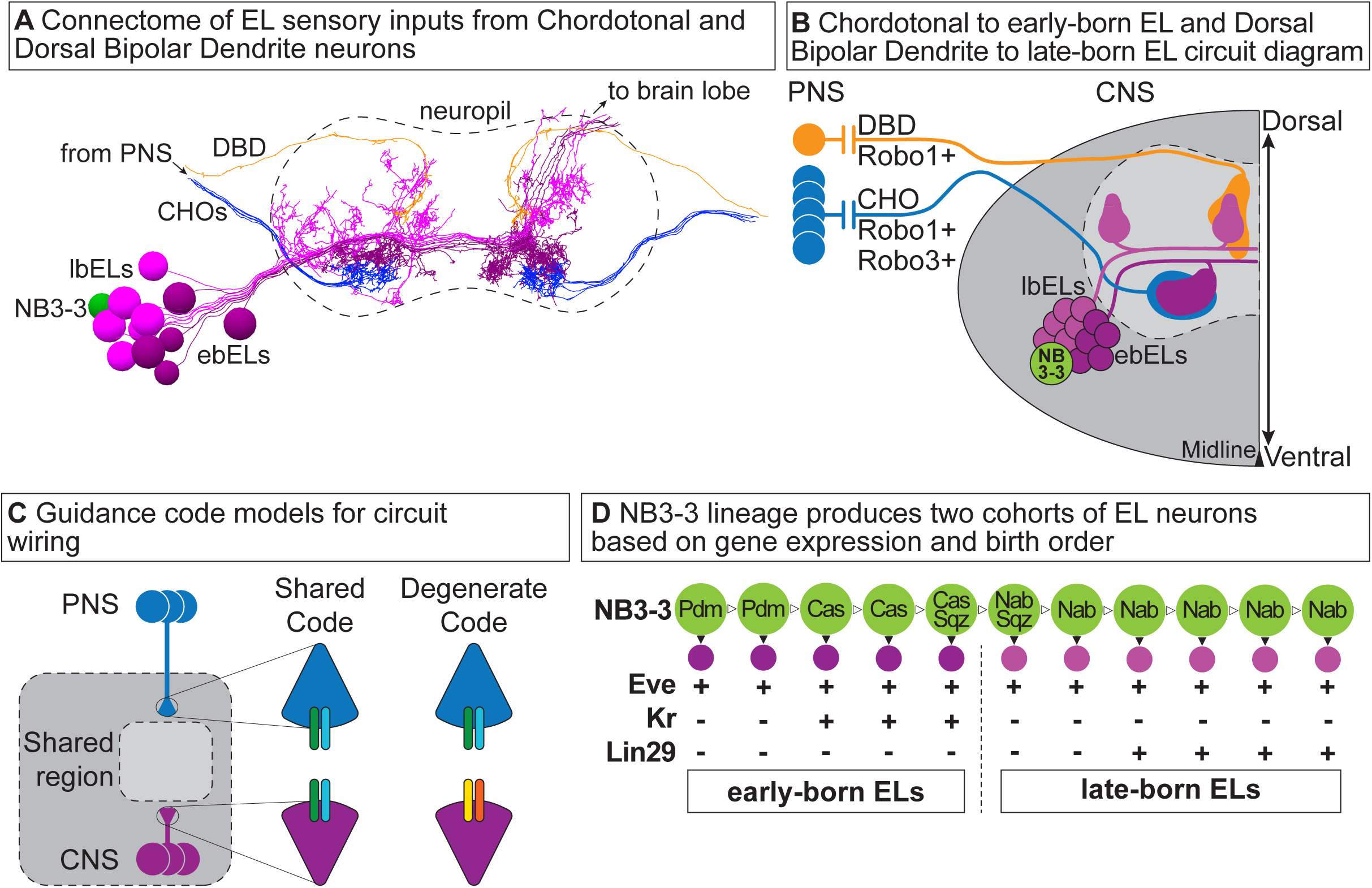
Circuit architecture of mechanosensory and proprioceptive circuits. **A** Image of wire skeleton of Chordotonal, blue, to early-born EL, dark purple, circuit and Dorsal Bipolar Dendrite, orange, to late-born EL, light purple, circuit from L1 larval CNS connectome shown in cross-section. Neuropil boarders represented by dashed outline. **B** Illustration of Chordotonal to early-born EL, and Dorsal Bipolar Dendrite to late-born EL circuit diagram. **C** Illustration of guidance cue models for circuit wiring. Sensory neurons, blue, and interneurons, purple, project to a shared region, light gray with dashed outline. Growth cones of neurites shown to express two potential codes of guidance expression. **D** Illustration of NB3-3 lineage. Green circle represents NB3-3, dark and light purple represent early-born and late-born ELs, respectively. Arrowheads represent terminal (filled in) and self-renewing (white) divisions. Each circle is one cell, and identity gene expression shown below: - represents no expression, **+** represents expression.

Our approach is to use the Drosophila embryonic and larval nerve cord to monitor the expression of Robo1 and Robo3 protein and mRNA. We find that, across the nerve cord, Robo3 is selectively transcribed in early-born neurons. Next, we use genetic manipulation of four Temporal Transcription Factors (TTFs)—Pdm, Castor, Sqz, and Nab to examine the regulation of *robo3* mRNA expression. We find that a subset of TTFs regulates *robo3* expression, and in some cases, this is genetically separable from TTFs’ regulation of classical temporal identity markers. Then, using cell-type-specific Robo3 perturbations, high-resolution morphological characterization, and calcium imaging, we evaluate the requirement for Robo3 in EL circuit anatomy and function. Within the EL circuit, Robo3 is required for dendrite positioning and branching, as well as for the normal high fidelity of vibrational sensory encoding. Together, our findings show that a shared guidance-receptor code links neuronal birth timing to circuit assembly.

## Results

The goal of this study was to better understand how axon guidance molecules are used in circuit assembly. To this end, we leveraged the well-defined stem cell lineages of the Drosophila embryonic nerve cord and Drosophila’s genetic toolkit to investigate the regulation of Roundabout 3 (Robo3) and its role in Even-skipped Lateral (EL) somatosensory circuit assembly. We sought to understand the pattern of Robo3 expression across the developing nerve cord, to identify genetic regulators of *robo3* transcription, and to define the role of Robo3 in circuit structure and function.

### Throughout the developing nerve cord, *robo3* is selectively transcribed in early-born neurons

To understand the role of Robo3 in circuit wiring, we needed to determine the extent to which it is differentially expressed in neurons of different birth times, including ELs of the NB3-3 lineage. We began our study by examining the distribution of Robo3 in the whole embryonic Drosophila nerve cord and in specific model lineages, and comparing it with the expression of Robo1, which is pan-neuronal^15,53^.

The Drosophila nerve cord is divided into an axon and dendrite-rich central neuropile surrounded by a soma-filled outer cortex. Roundabout receptor expression in the neuropile has been analyzed elsewhere^2,9,11,14,15,53–55^. Herein, we used anti-ElaV to visualize neural soma and examined the expression of Robo1 and Robo3 (Figure 2A-D). Robo1 protein puncta are evenly distributed in the cortex (Figure 2A’, B’), as expected for a pan-neural protein^15,53^. Robo3 puncta are near the neuropile, and absent from the lateral edges of the cortex (Figure 2C’, D’), demonstrating regulated expression. To determine whether Robo3 was regulated at the level of transcription, we generated *in situ* hybridization chain reaction probes^56^. Neither *robo1* nor *robo3* transcripts are detected in the neuropile (Figure 2A’’-D’’’). *robo1* transcripts are detected uniformly throughout the cortex (Figure 2A’’-B’’’). *Robo3* transcripts are detected medially but not laterally (Figure 2C’’-D’’’). Moreover, when visualizing *robo3* mRNA, we found heterogeneity in the levels (Figure 2E’’-G’’). In general, neurons expressing the highest levels of *robo3* mRNA expressed Robo3 protein (Figure 2E’-G’) and were located closer to the neuropile (Figure 2H-I). These data show that differential levels of Robo3 protein are at least partially controlled at the transcriptional level and *robo3* transcripts are enriched medially within the nerve cord.

**Figure 2.**
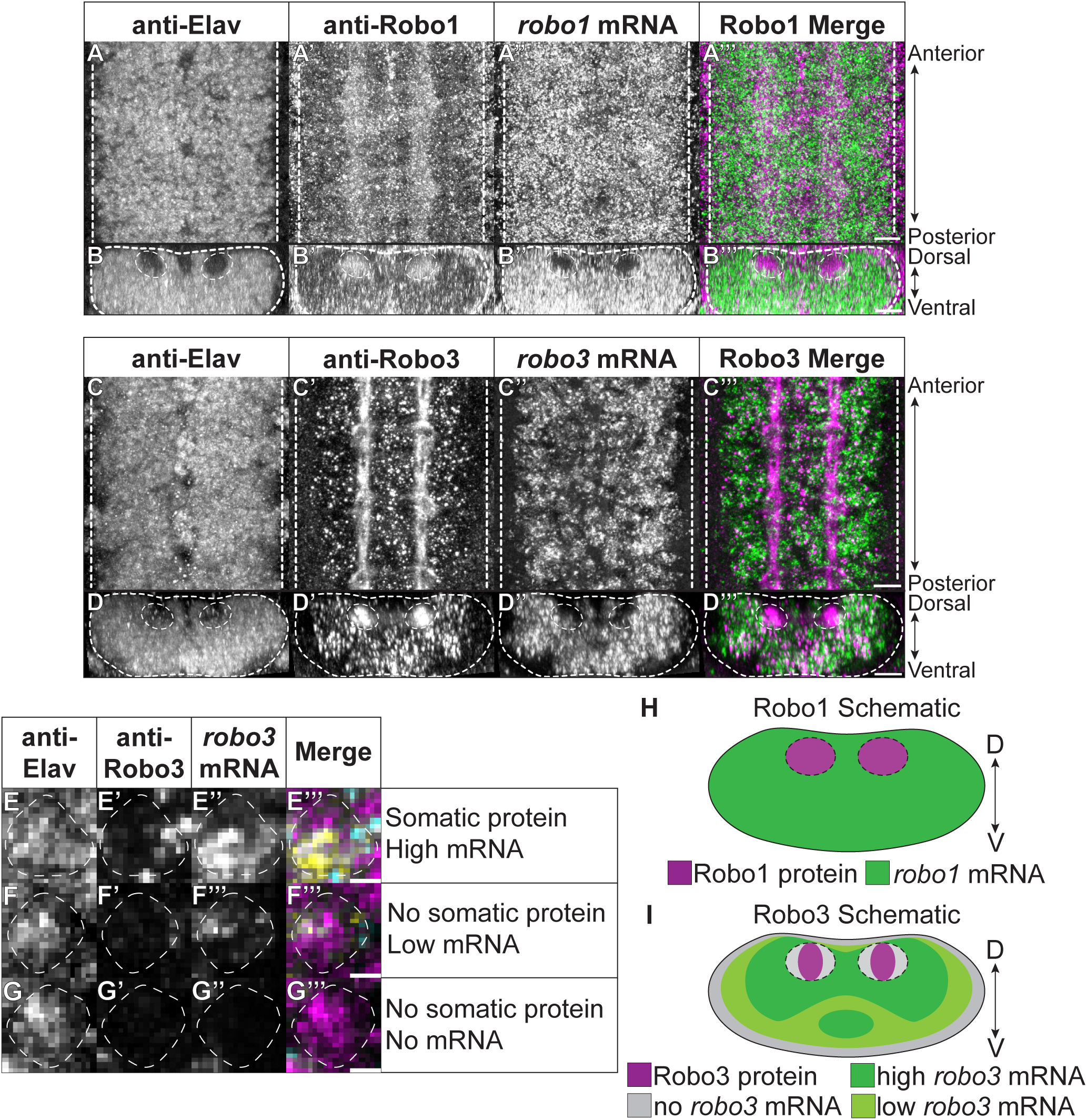
Robo3 mRNA and protein are strongly expressed in early-born neurons. **A-D** Images of Robo1 and Robo3 protein and mRNA expression in the Drosophila nerve cord of late-stage embryos. Dashed line denotes edge of CNS and neuropil. Scale bar represents 10 microns. **E-G** Images of the variance in Robo3 protein and mRNA expression at the single neuron level. The dashed line surrounds a singe cell. The scale bar represents 1 micron. **H-I** Illustration of Robo1 and Robo3 distribution in the wildtype embryonic nerve cord.

The medial enrichment of Robo3 protein and *robo3* mRNA suggests an association with neuronal birth time. In the Drosophila nerve cord, neural birth time and soma position are correlated, with soma of early-born neurons medial to those of later-born neurons^57^. So, we next co-labeled with early- and late-born markers. Kruppel (Kr) marks a subset of early-born neurons, and Lin29 marks most late-born neurons^50^. We used an anti-Kr antibody and generated a new probe for *lin29* mRNA. *Robo1* is expressed in all Kr-expressing and all Lin29-expressing cells (Supplemental Figure 1A-B, E-F). *robo3* is expressed in ∼one-half of Kr-expressing neurons, and ∼one-quarter of Lin29-expressing neurons (Supplemental Figure 1C, H). These data show *robo3* transcription is biased towards early-born neurons.

To more rigorously evaluate the relationship between Robo3 and neural birth timing, we used the NB3-3 lineage, in which EL birth timing is characterized at the single-neuron level. We stained with anti-Even-skipped (Eve), which stains the cluster of ∼10 ELs, and visualized Robo1 and Robo3 proteins (Figure 3A-B) or mRNA (Figure 3C-D). Robo1 protein puncta and mRNA were evenly distributed throughout the EL cluster (Figure 3A, Supplemental Figure 2). Robo3 puncta were enriched on the medial side of the EL cluster (Figure 3B). On average, eight ELs express *robo3* mRNA (Figure 3C-E). Three of three early-born Kr[+] ELs expressed *robo3* mRNA (Figure 3C, F), but only one of four late-born *lin29*[+] ELs expressed *robo3* mRNA (Figure 3D, G). We quantified the expression level of *robo3* in Kr[+] and *lin29*[+] ELs, finding significantly higher expression in early-born ELs (Figure 3H). Thus, early-born ELs express high levels of *robo3*, while the first late-born ELs express low levels of *robo3,* with the remaining late-born ELs not expressing *robo3* (Figure 3I).

**Figure 3.**
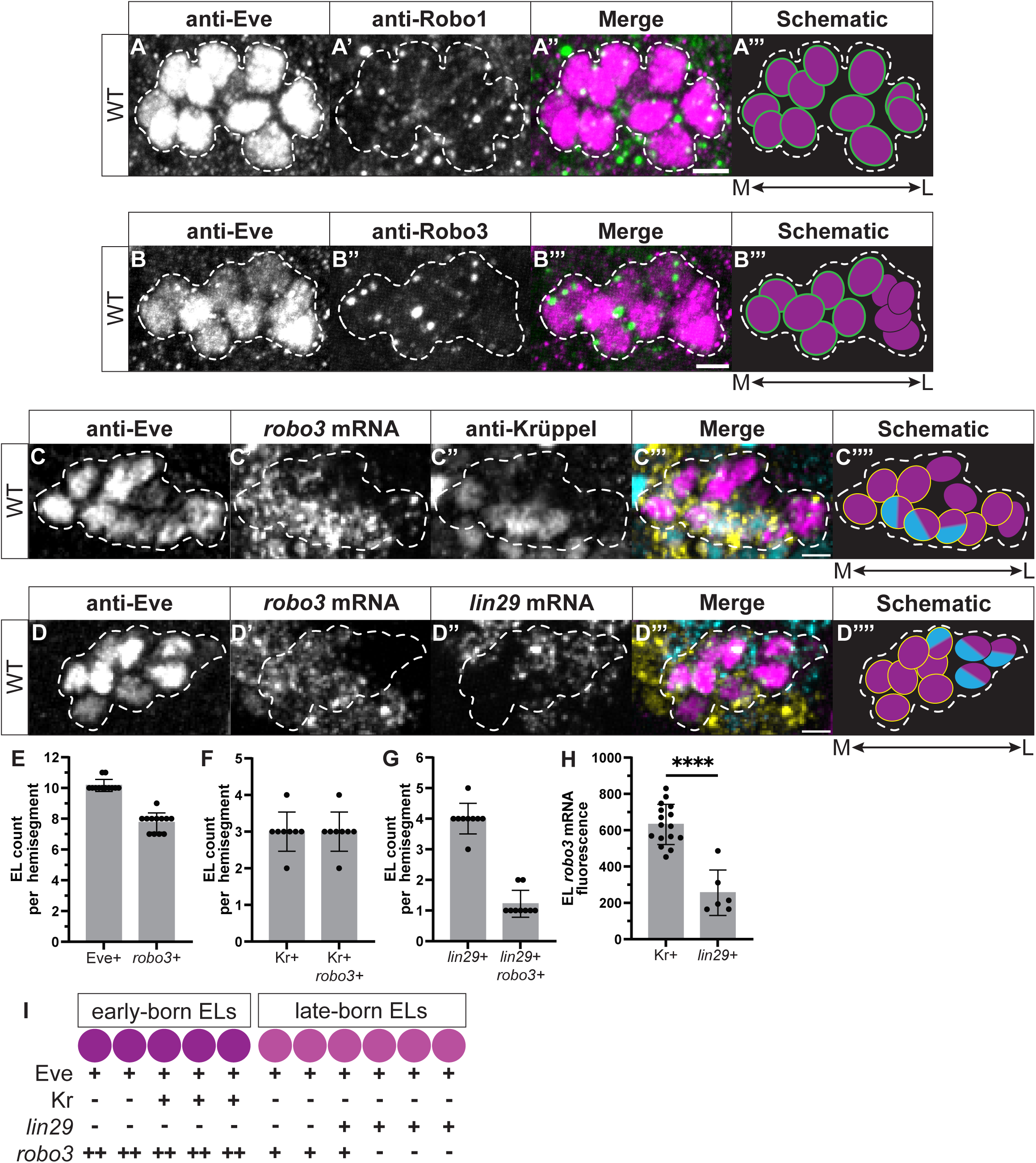
Robo3 protein and mRNA are expressed at high levels in early-born ELs, but not in the latest-born ELs. **A-D** Images and schematic summaries of ELs and Robo1 (A) and Robo3 (B) protein or *robo3* mRNA (C-D). EL cluster is outlined by dashes. Images show a single abdominal hemisegment from a late-stage embryo, with anterior up and midline to the left. Scale bars represent 2.5 microns. In schematics, each circle represents one cell, with Eve in magenta, guidance cue protein in green, mRNA in yellow, and temporal marker in blue. **E-H** Quantifications of *robo3* mRNA expression patterns. Each dot represents measurement in one hemisegment. Gray box shows average and black bars show standard deviation. Significance determined using Mann-Whitney test, **** = p<0.0001. **I** Illustration of *robo3* mRNA expression in the NB3-3 lineage. Each circle is one cell: - represents no expression, **+** represents expression, and **++** represents high expression.

To generalize our findings beyond one lineage, we also evaluated *robo3* mRNA expression in the NB7-1 lineage. Here, Eve is expressed in five U motor neurons, which are numbered according to birth order (i.e., U1 is the firstborn). *robo3* mRNA is expressed in U1-U4, but not U5 (Supplemental Figure 3A-G). We conclude that robo3 expression is correlated with U motor neuron birth order (Supplemental Figure 3O). Thus, across lineages, Robo3 transcripts are enriched in early-born neurons.

### Nab and Sqz regulate EL temporal identity but not *robo3* mRNA expression

Next, we sought to understand why *robo3* was highly expressed in early-born compared to late-born ELs. Regulators of temporal identity are prime candidates to control *robo3* mRNA expression because of their association with neural birth time and their ability to rewire circuitry in some lineages^36,37,58^. Thus, an appealing idea is that TTFs are upstream of the entire molecular program that directs circuit wiring, including axon guidance molecules. In NB3-3, a series of four temporal transcription factors (TTFs), Pdm, Cas, Sqz, and Nab, are sequentially expressed in the neuroblast as it gives rise to various ELs^50^. Of these TTFs, manipulation of Nab and Sqz alters EL temporal identity, as assessed by changes in the proportions of ELs expressing the Kr and Lin29^50^. Thus, in the NB3-3 lineage, we expected that TTFs regulate *robo3* expression downstream of their role in regulating temporal identity. We specifically hypothesized that Nab and Sqz act as negative regulators of *robo3* in ELs because they repress early-born EL identities and promote late-born EL identities (Figure 4A-B).

**Figure 4.**
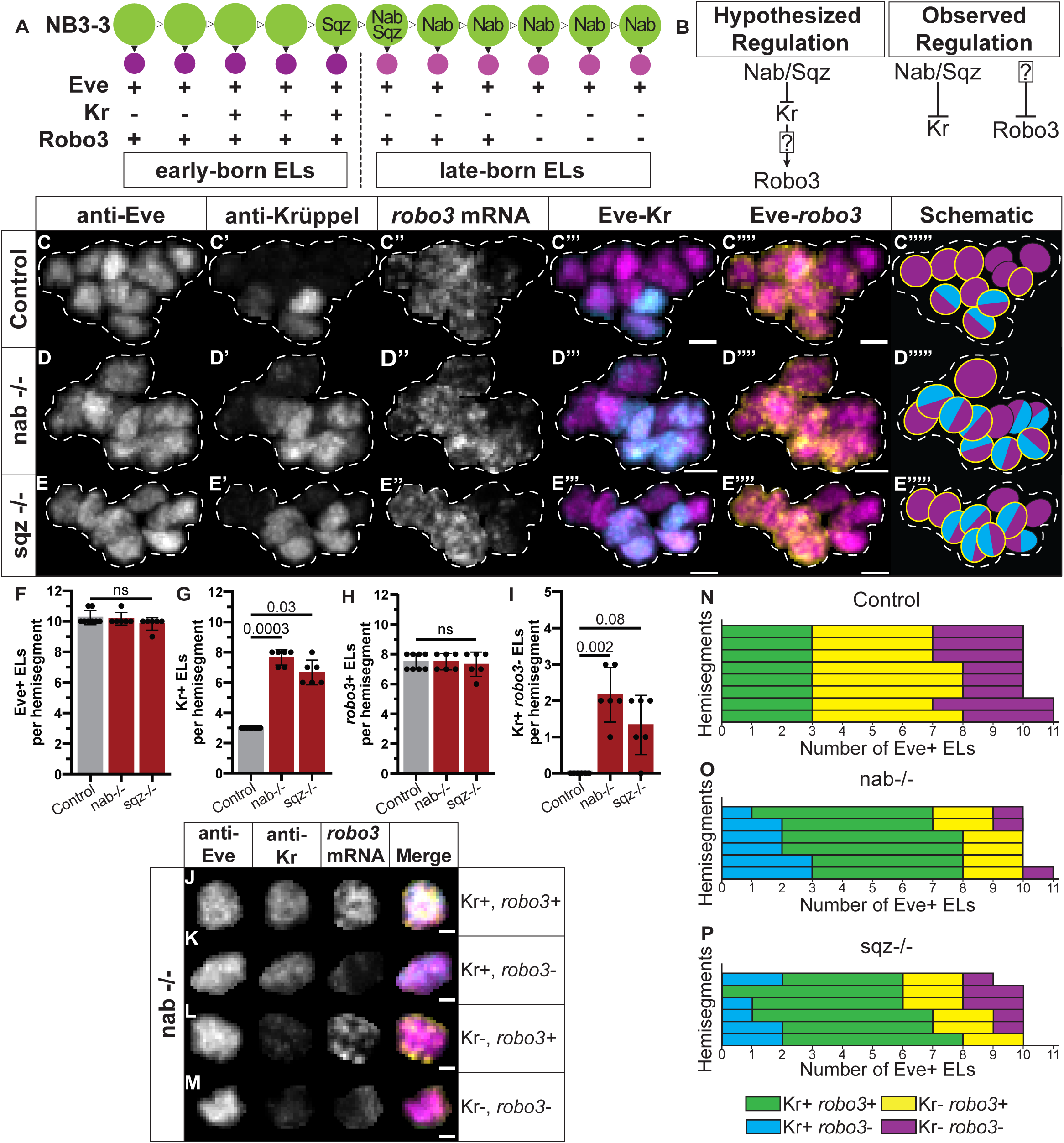
Nab and Sqz regulate early temporal identity but not *robo3* expression in ELs. **A** Illustration of NB3-3 lineage progression. Green circle represents NB3-3, dark and light purple represent early-born and late-born ELs, respectively. Arrowheads represent terminal (filled in) and self-renewing (white) divisions. -, +, ++ represent no expression, expression, and high expression. **B** Illustration of hypothesized and observed genetic relationships between Nab, Sqz, Kr, Lin29, and Robo3. **C-E** Images and schematic summaries of ELs, temporal identity, and *robo3* mRNA expression in ELs. EL cluster is outlined by dashes. Images who a single abdominal hemisegment from a late-stage embryo, with the anterior up and the midline left. Scale bars represent 2.5 microns. In schematics, each circle represents one cell, with Eve in magenta, the *robo3* mRNA in yellow, and Kr temporal marker in cyan. **F-I** Quantifications of expression patterns. Each dot represents the measurement in one hemisegment. Gray box shows average, and black bars show standard deviation. Significance determined using Kruskal-Wallis test (F p>0.05, G p<0.0001, H p>0.05, I p<0.005) and Dunn’s multiple comparisons (ns = not significant, p-values reported above each comparison). **J-M** Images of Kr and *robo3* expression at single EL level. Merge represents Eve in magenta, Kr in cyan, and *robo3* in yellow. Scale bars represent 1 micron. **N-P** Quantifications of Kr and *robo3* co-expression in ELs Each row represents a different hemisegment. Genotypes: control is nab^G26^ heterozygous ; nab -/- is nab^G26^ homozygous, sqz -/- is sqz^02102^ homozygous.

First, we confirmed Nab’s role in regulating EL temporal identity. In both control and *nab* mutants, the number of ELs in the lineage was unchanged (Figure 4C-D’’’’’, F). In the *nab* mutant, the number of Kr[+] ELs more than doubled (Figure 4C’-D’’’’’, G), whereas *lin29*[+] ELs were completely lost (Supplemental Figure 4). Therefore, Nab regulates temporal identity in ELs, consistent with prior findings.

Next, we determined the role of Nab in regulating *robo3* expression. Surprisingly, in *nab* mutants, the number of *robo3*[+] ELs remained the same (Figure 4C”-D’’’’’, H). We quantified the co-expression of Kr and Robo3 (Figure 4I-L). In *nab* mutants, we found 1 to 3 Kr[+] *robo3*[-] ELs, which was never seen in controls (Figure 4J, M-O). Thus, *robo3* and temporal marker gene expression were genetically separable in the *nab* mutants. We conclude that in ELs, neither Nab nor Lin29 represses *robo3* expression, and Kr does not promote *robo3* expression.

To determine if this unanticipated phenotype was specific to Nab, we performed a similar analysis in *sqz* mutants. In *sqz* mutants, the number of Kr[+] ELs also increases without changes to *robo3* expression (Figure 4E-H), and Kr[+] and *robo3[-]* ELs are found in the *sqz* mutants (Figure 4M,P). Thus, Sqz also does not regulate *robo3* in ELs.

In summary, our results show that in the ELs of the NB3-3 lineage, *robo3* expression is not regulated by Nab, Sqz, Lin29, or Kr (Figure 4Q). Thus, contrary to our expectations, the temporally associated pattern of *robo3* expression cannot be explained by a simple model in which TTFs act as master regulators of circuit assembly. Instead, we find that the regulation of neural temporal identity and guidance receptors is genetically separable.

### Pdm and Castor regulate temporal identity and *robo3* expression in multiple lineages

We remained curious about the strong association between *robo3* expression and neural birth order (Figures 1, 2). We wanted to be comprehensive in our analysis of the proposed NB3-3 TTFs and, therefore, assayed Pdm and Cas.

Despite well-defined roles in other lineages the roles of Pdm and Cas in regulating temporal identity in the NB3-3 lineage have not yet been determined^59–61^. We used the early-born marker, Kr, to assay changes in temporal identity in *pdm* and *cas* mutants. Pdm is expressed in NB3-3 as it divides to give rise to the first- and second-born ELs, and Cas is expressed in NB3-3 as it divides to make the Kr[+] third-to-fifth-born ELs^50^ (Figure 5A). In *pdm* mutants, the number of ELs is reduced by one to two (Figure 5B-C, E), consistent with the idea that NB3-3 skips the production of the first- and second-born ELs. The number of Kr [+] ELs was reduced by one (Figure 5B’-C’, F), indicating that Pdm has effects on the identity of ELs. In *cas* mutants, the number of ELs is unchanged compared to controls (Figure 5D-E), suggesting shifts rather than skips in EL temporal identities. The number of Kr[+] ELs was reduced by two to three (Figure 5D’, F), showing that third-to-fifth born EL identities are altered. We conclude that Pdm influences EL numbers and that Pdm and Cas influence the temporal marker Kr in the NB3-3 lineage.

**Figure 5.**
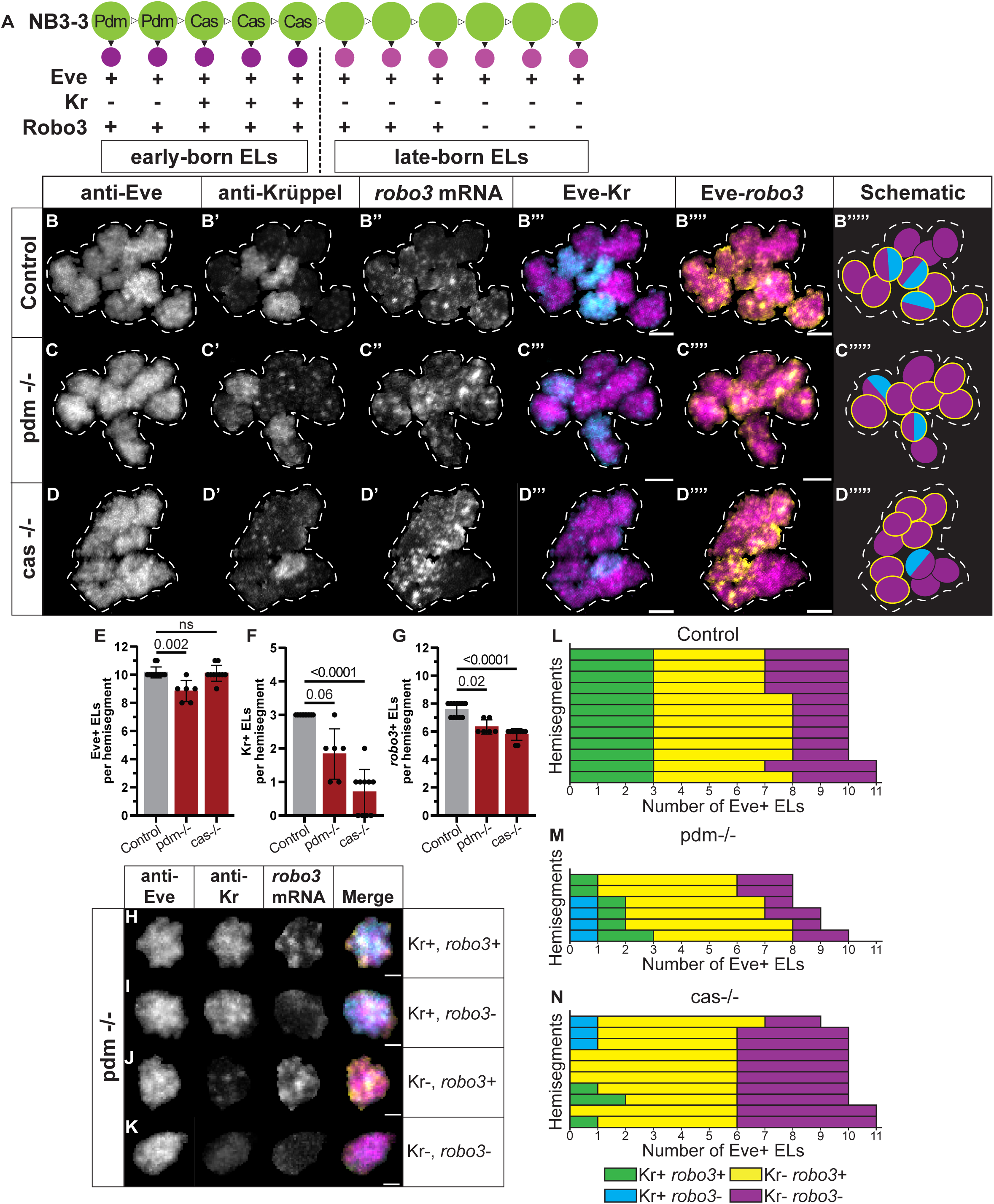
Pdm and Cas regulate early temporal identity and *robo3* expression in ELs. **A** Illustration of NB3-3 lineage progression. Green circle represents NB3-3, dark and light purple represent early-born and late-born ELs, respectively. Arrowheads represent terminal (filled in) and self-renewing (white) divisions. -, +, ++ represent no rexpression, expression, and high expression. **B-D** Images and schematic summaries of ELs, temporal identity, and *robo3* mRNA expression in ELs. EL cluster is outlined by dashes. Images who a single abdominal hemisegment from a late-stage embryo, with the anterior up and the midline left. Scale bars represent 2.5 microns. In schematics, each circle represents one cell, with Eve in magenta, the *robo3* mRNA in yellow, and Kr temporal marker in cyan. **E-G** Quantifications of expression patterns. Each dot represents the measurement in one hemisegment. Gray box shows average, and black bars show standard deviation. Significance determined using Kruskal-Wallis test (E p=0.002, F p<0.0001, G p<0.0001) and Dunn’s multiple comparisons (ns = not significant, p-values reported above each comparison). **H-K** Images of Kr and *robo3* expression at single EL level. Merge represents Eve in magenta, Kr in cyan, and *robo3* in yellow. Scale bars represent 1 micron. **L-N** Quantifications of Kr and *robo3* co-expression in ELs Each row represents a different hemisegment. Genotypes: control is cas^j1c2^ heterozygous, pdm -/- is Df(2L)ED773 homozygous, cas -/- is cas^j1c2^ homozygous.

We next evaluated *robo3* expression. In *pdm* and *cas* mutants, the number of *robo3*[+] ELs was reduced by two in comparison to controls (Figure 5B’’-D’’’’’, G), indicating both transcription factors are partially required for *robo3* expression. In both mutants, a subset of Kr[+] ELs lack *robo3* expression (Figure 5H-K). Thus, Pdm and Cas regulate Kr expression and *robo3* expression in ELs in a manner that is not causally linked.

To determine whether *cas* regulated *robo3* in other lineages, we used the U MNs in NB7-1. *cas* mutants have three Kr[+] U MNs and multiple Kr[-] U4s, as expected^60^ (Supplemental Figure 3A-K). In *cas* mutants, all Kr[+] U MNs express *robo3* (Supplemental Figure 3F-H, O-P), and the majority U4s express *robo3* (Supplemental Figure 3I, L, N). Thus, in NB7-1, *robo3* expression changes correlate with alterations in temporal patterning (Supplemental Figure 3O-P).

In summary, our data demonstrate that in NB3-3, Pdm and Cas regulate Kr and *robo3* expression, but their effects are separable. In the NB7-1 lineage, Cas regulates *robo3* in concordance with its regulation of temporal identity. Most generally, we find that across lineages, Cas regulates *robo3*, although the details are lineage-specific.

### Roundabout 3 is required in early-born ELs for dendrite positioning and branching

Our investigation of Robo3 in ELs was motivated by the observation that for ELs’ sensory neuron inputs (Figure 1?), differences in Robo3 play a key role in positioning sensory neuron dendrites. Specifically, CHOs express Robo3 whereas DBDs do not (Figure 1B)^52^.

Because CHOs synapse with early-born ELs that express high levels of Robo3, whereas DBDs synapse with late-born ELs that express little to no Robo3 (Figure 2) suggests a model in which neurons of different origins extend their neurites to a shared location within the CNS through the use of a shared code (Figure 1C). Further, these data argue against the use of a degenerate code, in which each neuron employs a unique combination of guidance receptors (Figure 1C).

However, to rigorously support a shared code model and reject a degenerate code model, Robo3 must be required in ELs for normal circuit architecture and function.

Because Robo3 is highly expressed in many early-born neurons in addition to early-born ELs (Supplemental Figure 1), we needed to investigate its cell-autonomous role in regulating neural morphology. To selectively disrupt Robo3 function in ELs, we obtained a UAS-fly line with an intact extracellular portion and replaced the cytoplasmic portion with Myc tags, here termed “*UAS-Robo3-ΔC*” (Figure 6A) to generate a dominant-negative as has been done for other Robo proteins^54,55,62–64^.

**Figure 6.**
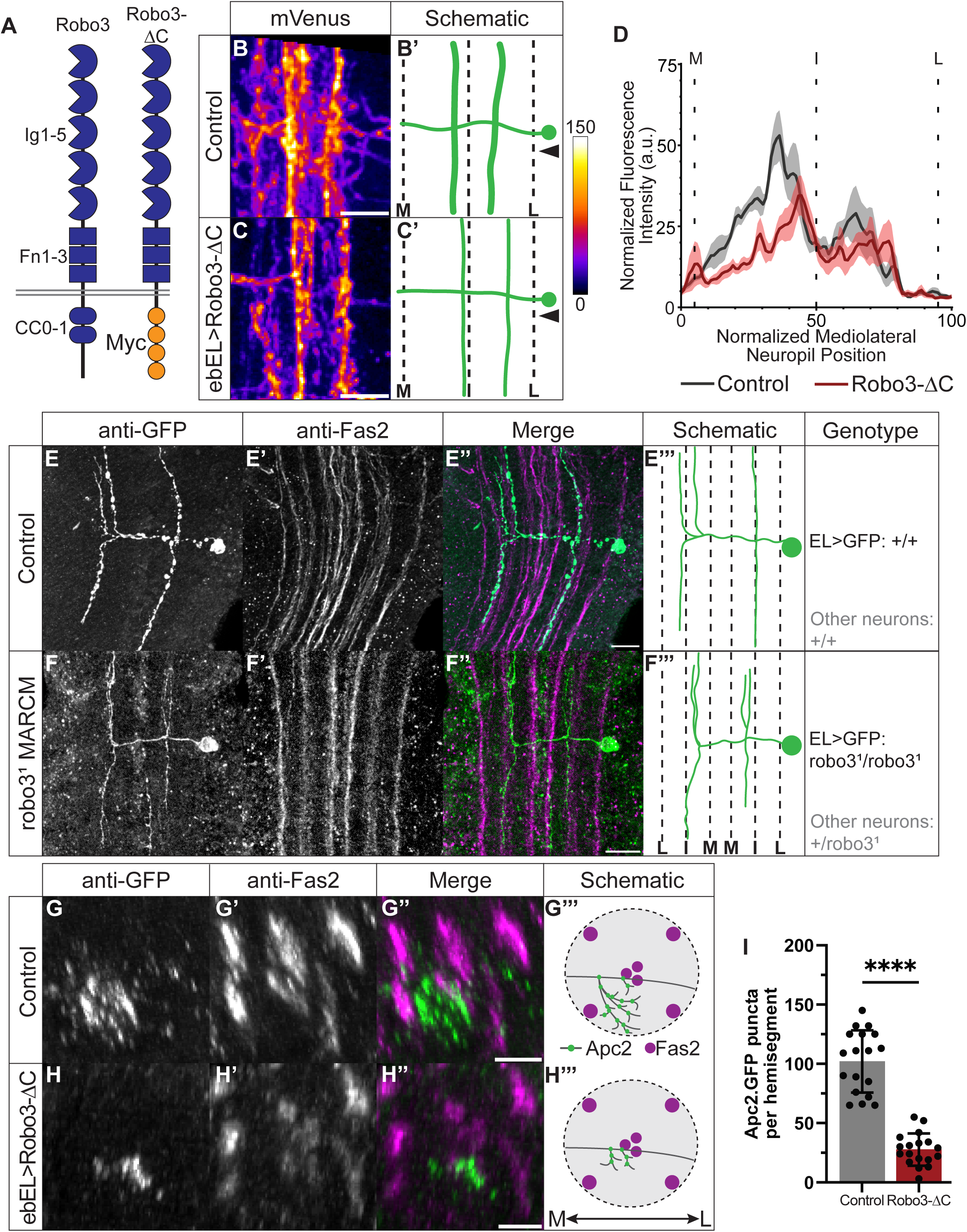
Early-born ELs require Robo3 for dendrite positioning and arborization. **A** Illustration of Robo3-ΔC protein structure in comparison to wildtype Robo3. **B, C** Images and schematic summaries of control and Robo3-ΔC early-born ELs expressing mVenus. Green represents GFP, dashed lines represent Fas2. Arrowhead represents line scan quantification. Medial (M), intermediate (I), and lateral (L), fascicles are labeled. **D** Quantification from line-scan of membrane fluorescence across abdominal hemisegment. **E, F** Images and schematic summaries of control and *robo3*^1^ homozygous early-born EL single cell clones. **G, H** Images and schematic summaries of control and Robo3-ΔC early-born ELs expressing Apc2.GFP. **I** Quantification of Apc2.GFP puncta of early-born ELs. Each dot represents the measurement of one hemisegment. Boxes show average and black bars show standard deviation. Significance determined using Welch’s t-test, **** = P<0.0001. All images are from 1^st^ instar larval VNC dissections. B-F are shown from dorsal view, G and H are shown in cross-section. Scale bars represent 10 microns. Genotypes: control is WT robo3 homozygous, in robo3^1^ MARCM ELs are robo3^1^ homozygous while the surrounding tissue is robo3^1^ heterozygous.

We assayed early-born EL morphology in larvae after the circuits had matured. Using the *early-born-EL-GAL4* line, we expressed *UAS-membrane-GFP* with or without *UAS-Robo3-ΔC*. This method allows us to visualize all the neurites of early-born ELs, which form complex patterns because typical ELs project across the midline and across segments^31,41,51,65^. To simplify our analysis, we focused on a portion of the arborization within a single left or right half segment (hemisegment) of the first abdominal segment (A1), pseudo-coloring the GFP channel to visualize the most prominent arbors (Figure 6B, C). To get a more quantitative description, we measured and plotted GFP fluorescence intensity along a line drawn from medial to lateral (Figure 6B-D). Also in these experiments, we co-stained with Fas2 to provide landmarks within the neuropile^66^. In controls, early-born ELs project to the anterior and posterior in two thick longitudinal bundles, with less prominent arborizations enriched towards the midline (Figure 6B,D). The bundles were located on either side of the Fas2-labeled intermediate fascicle, which appeared as two fluorescence peaks along the medio-lateral axis (Figure 6B,D). When Robo3 activity is disrupted in early-born ELs, longitudinal projections and medial elaborations are both reduced (Figure 6B-D). These data suggest Robo3 activity is required in early-born ELs for their normal morphology. However, this visualization is quite coarse.

To obtain a higher-resolution understanding of the role of Robo3 in ELs, we made single-neuron *robo3* null mutant clones using Mosaic Analysis with a Repressible Cell Marker (MARCM) (Figure 6E)^67^. We restricted our analyses to neurons in segment A1 because the morphologies of the ELs in that segment are well characterized^41^. In wild-type, a majority of early-born ELs in A1 have an “H-shaped” morphology (Figure 6F, F’’’): Two dendritic branches project anterior-posterior on either side of the midline just lateral to the intermediate fascicle. An axon projects to the anterior just medial to the intermediate fascicle such that the axon and dendrite are found on either side of the intermediate fascicle and are separated from each other (Figure 6F’’, F’’’). In a *robo3* null mutant clone, the overall H-shaped morphology is preserved (Figure 6G). However, the separation between the dendrite and axon branches is lost because the dendrites shift medially, but the axon does not (Figure 6G’’, G’’’). Thus, Robo3 normally repels early-born EL dendrites from the midline. Two caveats to these experiments are that we were only able to generate a few clones in segment A1, so we lacked sufficient replicates to quantify the effect. Furthermore, MARCM labeling enabled visualization of the EL’s major branches, but not the finer dendritic branching.

Next, we assayed the role of Robo3 in early-born EL dendrite branching. Again, we used *early-born-EL-GAL4,* with or without *UAS-Robo3-ΔC,* this time with *UAS-Apc2.GFP* to mark the location of microtubule branch points found at dendritic branches^68^. The Apc2.GFP forms foci that are clearer when observed in cross-section (Figure 6H, I). In control, A1 hemisegments have ∼100 Apc2.GFP puncta (Figure 6H, J), most positioned medially and ventrally to the intermediate Fas2 fascicle (Figure 6H’’, H’’’). When Robo3 activity is disrupted in early-born ELs, the number of branch points was reduced by 80% (Figure 6J), and the puncta occupied a small domain just ventral to the intermediate Fas2 fascicle (Figure 6I’’, I’’’). Thus, ELs require Robo3 for proper branching.

We conclude that Robo3 activity is required cell-autonomously in early-born ELs to repel dendrites from the middle and to promote dendrite branching.

### Roundabout 3 is required for the fidelity of early-born EL vibration response

In our final set of experiments, we examined the role of Robo3 in early-born EL function. While the effects on morphology could translate to functional defects, recent studies have shown that axon guidance molecules can be dispensable for normal function due to CNS plasticity^69,70^. CHO sensory neurons and early-born ELs both respond to vibrational stimuli: Specifically, CHOs respond to the entirety of a long vibrational stimulus, and early-born ELs quickly respond with a fast decay even as the vibration persists (Figure 7A)^41,65^. We need to understand how early-born ELs lacking normal Robo3 function respond to vibration.

**Figure 7.**
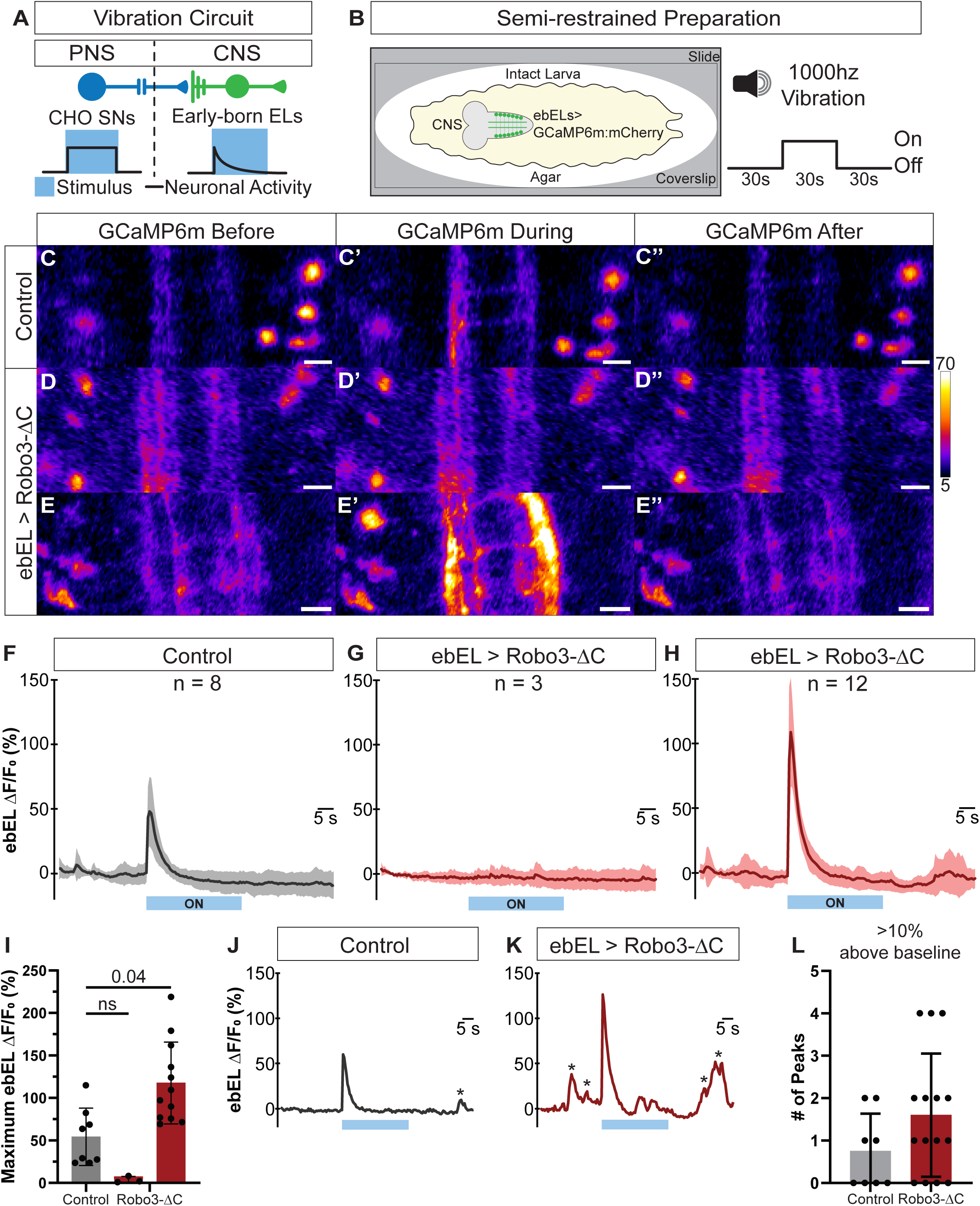
Early-born ELs require Robo3 for faithful responses to vibrational stimuli. (A) **A** Illustration of CHO-to-ebEL vibration circuit with activity below each neuron class. CHO sensory neuron in dark blue, early-born EL in purple. **B** Illustration of semi-restrained preparation for live images of intact larvae. Neuronal activity recorded for 90 seconds (s), with a 30s a 1000 hertz sound stimulus played in the middle. **C-E** Images from control and Robo3-ΔC early-born ELs expressing GCaMP6m live calcium imaging trials, with one frame from before, during and after stimulus from intact L1 larvae. Scale bars represent 10 microns. **F-H** Quantification of normalized change in GCaMP fluorescence over mCherry fluorescence in control and Robo3-ΔC larvae. Robo3-ΔC split between two phenotypic classes. Average and standard deviation represented through dark line and lighter ribbon respectively. Stimulus on during light blue bar. **I** Quantification of max change in normalized fluorescence. Each dot represents one larval recording. Box shows average and black bars show standard deviation. Significance determined using Kruskal-Wallis test (p = 0.001) and Dunn’s multiple comparisons (ns = not significant, p-value reported above comparison). **J-K** Single recording of non-stimulus associated deflections from single larval recording. **L** Quantification non-stimulus associated deflections. Each dot represents number of peaks observed per larvae.

We used a calcium imaging assay where intact larvae are semi-immobilized between an agarose pad and a glass coverslip (Figure 7B)^71^. Larvae can perform small movements but not translocate out of the imaging window. We used *early-born-GAL4* with or without *UAS-Robo3-ΔC* to drive *UAS-GCaMP6m.mCherry.* Both fluorescent signals were simultaneously recorded before, during, and after a 30-second-long 1000 Hz stimulus (Figure 7B). The mCherry signal controlled movement artifacts and the GCaMP signal reported neural activity (Supplemental Figure 5). After the recording, we allowed larvae to recover and discarded recordings from immotile larvae to control for animal health.

In controls, as expected, at the onset of vibration, early-born ELs robustly responded with a quick decay (Figure 7C-C’’, F, I). Upon disruption of Robo3 activity in early-born ELs, we observed two phenotypes: In the minority (n=3 preparations), ELs did not respond, indicating that vibration encoding was eliminated (Figure 7D-D’’, G, I). In the majority (n=12 preparations), early-born ELs responded at the onset of vibration, with a marked increase in GCaMP intensity (Figure 7E-E’’, H, I). We quantified the peak change in GCaMP fluorescence intensity relative to the mCherry signal, finding a 2-fold increase in GCaMP fluorescence when Robo3 was disrupted in early-born ELs compared with controls (Figure 7I). In this set of preparations, we noticed a second phenotype: many small spontaneous firing events occurring before and after the stimulus (Figure 7J-K). We quantified the number of spontaneous firing events that exceeded 10% above baseline and observed an overall increase in early-born ELs in Robo3-disrupted animals compared with controls (Figure 7L). Thus, without normal Robo3 function, early-born-EL encoding of vibrational stimuli is highly variable, but never normal.

In summary, perturbation of Robo3 in early-born ELs reduces the fidelity of their vibrational response. We conclude that proper Robo3 function in early-born ELs is critical for their normal circuit-level somatosensory encoding.

## Discussion

In this study, we sought to understand the relationship between neuronal birth order and circuit assembly by examining regulation and necessity of Robo3 during somatosensory circuit wiring. This study spans levels of analysis –starting with a well-studied axon guidance receptor, identifying a new developmental pattern of expression, its transcriptional regulation, and a new role at the cell level regulating dendritic morphology and circuit function. Previously, Robo3 was shown to be required in chordotonal sensory neurons which directly synapse onto early-born ELs within the ventral-intermediate region of the neuropil (Figure 1). Like their sensory inputs early-born ELs express high levels of Robo3 and that Robo3 expression is generally enriched in Kr[+] early born neurons across the VNC (Figure 2, 3, SFigure 1). To understand the Robo3 pattern we examined the temporal transcription factors: Nab, Sqz, Pdm, and Cas. We confirm the negative regulation of Kr by Sqz and Nab, while newly showing that Pdm and Cas positively regulate Kr in ELs (Figure 4, 5). We find that only Pdm and Cas can regulate Robo3 expression in ELs (Figure 4, 5). Robo3 in early-born ELs is necessary for their medio-lateral dendrite positioning and dendritic branching (Figure 6). Robo3 in early-born ELs is required for their normal encoding of vibrational stimuli (Figure 7). These insights propose a shared, rather than divergent, guidance receptor code (Figure 1) connect neural birth timing to circuit assembly.

## Robo3 is enriched in early-born neurons

How guidance is regulated at the mRNA and protein levels is a critical and lingering question in researching neuronal circuit wiring^21^. Many transcription factors have been identified to regulate guidance receptor expression, including Atonal, Eve, Hb9, Islet, Lhx, and Nkx2.9^17,44,46–49,52,72^. This research indicates that when neuronal identity is specified guidance receptor expression is conferred. However, transcription factor regulation of static neuronal identity does not clarify how guidance is regulated temporally. Previous work showed that Robo3 protein expression appears later during Drosophila development in comparison to Robo1 and 2 protein^15^. Altogether this could indicate that as neurons are born later in development they express Robo3. To understand the Robo3 expression pattern we needed to interrogate with lineage and temporal identity markers at a single cell level. We newly show that at the tissue and lineage levels early-born neurons are turning on *robo3* hours after they differentiate (Figure 2, 3). This demonstrates a relationship between neuronal birth order and guidance receptor expression. This leads us to question to what extent is this principle generalized? To what extent is the entire Robo code—robo1, robo2, and robo3—birth order regulated. For example, does *robo2* display a similar or opposite pattern to *robo3*? Are other mediolateral receptors, for Netrin, Fra and Unc-5 expressed in association with neuronal birth order? More experiments are needed to understand whether Robo3 is uniquely enriched in early-born neurons, or if guidance receptors fall into different classes of transcriptional regulation. With this new insight future research can elaborate on the relationship between temporal regulation and guidance cues.

### Temporal transcription factor regulation of identity and Robo3 expression is correlated but not coupled with neuronal birth time

Given the strong association of *robo3* expression with birth time, our data suggested that *robo3* expression could be controlled by regulators of temporal cell fate. Several studies have suggested that TTFs are upstream of circuit wiring, which would indicate that they are potential upstream regulators of guidance receptors^36,37,39,58,73^. According to this model, manipulation of any TTF that changes temporal fate, should alter *robo3* expression^30,50^. We found this to be true in the NB7-1 lineage, with the expansion of *robo3*[+] U4-like neurons in whole embryo cas mutants. However, in the NB3-3 lineage there is more complex relationship between TTFs and *robo3* expression. It was previously reported that nab and sqz are regulators of late-born identity in NB3-3 (Figure 4)^50^. In NB3-3, we newly show that Pdm and Cas also act as regulators of temporal identity (Figure 5). It was possible that each TTF regulated *robo3* expression downstream of temporal birth order markers like Kr which are know to be potent terminal selectors^74^. Unexpectedly, Nab and Sqz fail to regulate *robo3* expression and only Pdm and Cas act as positive regulators of *robo3* in ELs. It remains unclear what may repress *robo3* expression in the subset of late-born ELs.

From these results we propose two potential timing mechanisms for the regulation of temporal identity and *robo3* expression in the NB3-3 lineage. The first mechanism is a purely lineage intrinsic one in which TTFs have a modular capacity to regulate gene expression during the specification of neural progeny. In the case of the NB3-3 and ELs, Pdm and Cas regulate both Kr and *robo3* expression with some variance in affinity, while Nab and Sqz only regulate Kr. Due to the co-expression of identity regulators or other transcription factors each TTF produces different transcriptional outcomes. The second mechanism is a combined intrinsic-extrinsic mechanism in which timing is set within the lineage by TTFs and feedback from the surrounding environment impacts gene expression within neural progeny. This feedback can originate from any non-cell autonomous source like other neurons, glia, or secreted cues within the extracellular environment. In this scenario for NB3-3 and the ELs each TTF sets the timing of temporal identity markers like Kr and signals from the environment regulate *robo3* expression in early-born ELs. We consider both mechanisms to be plausible because our TTF manipulation are whole animal mutations. Therefore, we are not only affecting TTF expression within NB3-3 but all other cells within the CNS as well. Similar timing mechanisms have been proposed for protein regulation in the visual system and commissural axons^75–77^. To evaluate the strength of either temporal mechanism, future research will need more finely controlled or mosaic manipulations of TTFs within model lineages. Overall, it is critical to understand what connects TTF regulation of birth timing with Robo3 for circuit assembly, and how that relationship works alongside known terminal selector expression^50,73,78^.

### In interneurons, Robo3 is required for dendrite positioning and branching, and sensory processing function

We expected to find that *robo3* regulates EL dendrite position since its role in dendritogenesis has been recently appreciated as feature of neuronal guidance^79–82^. We newly show that Robo3 is critical for not only position, but dendrite arborization. Our data show that Robo3 is necessary to establish stabilized microtubule branches that give dendrites their mature morphology (Figure 6). Likely this is due to the role of guidance cues in regulating growth via the cytoskeletal network^20,83,84^. Robo1 is known to regulate microtubule assembly via the tyrosine kinase Abl and CLASP^54,83^. Yet, how Robo3 specifically regulates cytoskeletal rearrangements like microtubule branch points for dendrite morphogenesis has not been evaluated. Robo3 processes Slit binding differently than Robo1 due to the different cytoplasmic domains which are critical for longitudinal paths selection^55^. However, this does not fully clarify how the complex dendritic morphology of interneurons is achieved from Robo3 signaling.

Interaction between Robo3, Robo1, and other guidance cues may determine what morphology is ultimately stabilized during development. Further studies are needed to understand how Robo3 regulates cytoskeletal rearrangements and the establishment of dendritic morphology.

The role of Robo3 is not only important for morphology of early-born ELs but also their function. Previously, it was reported that when Chordotonal sensory axons are forced to terminate in an aberrant location their downstream interneurons can overcome the significant shifts in position and still establish connections^70^. Therefore, it was concluded that downstream interneurons rely on partner derived cues to establish functional connections^70^. However, our data show that guidance information from Robo3 is necessary in downstream sensory processing ELs for their faithful response to sensory stimuli (Figure 7). In the both our data and the previous research, downstream sensory neurons displayed stronger evoked responses following stimulation or activation of the sensory neurons. Due to these similar phenotypes, we conclude that guidance cues are required in both the sensory and downstream interneurons to refine the functional connections between circuit partners. When spatial positioning is disrupted, interneurons are still able to establish synapses with their circuit partners, yet that the percentage of sensory inputs on to those interneurons has changed (Figure 7)^70^. This indicates that guidance cues like Robo3 are not only important for neuronal pathfinding but a necessary and potentially modulatory step to achieve a specific circuit function. Future work will need to investigate the relationship between guidance cues and circuit establishment in interneurons to determine the if they play a direct role in establishing synaptic connections.

### A shared Robo code is used by synaptic partners from different developmental origins

Due to the different origins and environments that sensory and interneurons have to grow through to establish synaptic connections it was possible that they use unique guidance codes to find the same spatial regions. Our data reveals, however, that neurons from different developmental origins (e.g., PNS and CNS) send their neurites to a shared location with the CNS using a shared code of guidance cues. A shared guidance receptors code has been observed previously where Sema2b positive neurons arrived along the same longitudinal tract^85^. While Sem2b signaling is critical for larval response to vibrational stimuli it was unclear the circuit member of all Sema2b[+] neurons. Additionally, the authors indicate that the lateral position of Sema2b[+] neurons takes advantage of Robo3 signaling. In our study, we take advantage of the depth of knowledge about the somatosensory circuits and show that both chordotonals and early-born ELs require Robo3 to establish a functional circuit together. Together both ours and previous research demonstrate that a shared guidance is critical for the circuit assembly of neurons from different developmental origins. Chordotonals and early-born ELs only represent two members of their larger circuit architectures that make up the mechanosensory and proprioceptive processing circuits^31^. Future work will be needed to fully evaluate how guidance is regulated across multiple synaptic partners, and determine if a shared code is always necessary for circuit wiring

### Conclusions and Outlook

We identify at the transcriptional level how a shared Robo3 code is established in early-born neurons via the regulation of temporal transcription factors. Additionally, we demonstrate how this code is required for both neuronal morphology and function. We only evaluate *robo3* expression and regulation in two lineages within the Drosophila VNC abdominal segments. In other lineages and regions of the CNS we may observe these mechanisms adjusted to the different spatial and temporal contexts^41,86^. Our research provides insights into a new mechanism for how neuronal birth timing regulates circuit assembly through the TTF regulation of a shared guidance receptor code.

### Key Resources Table

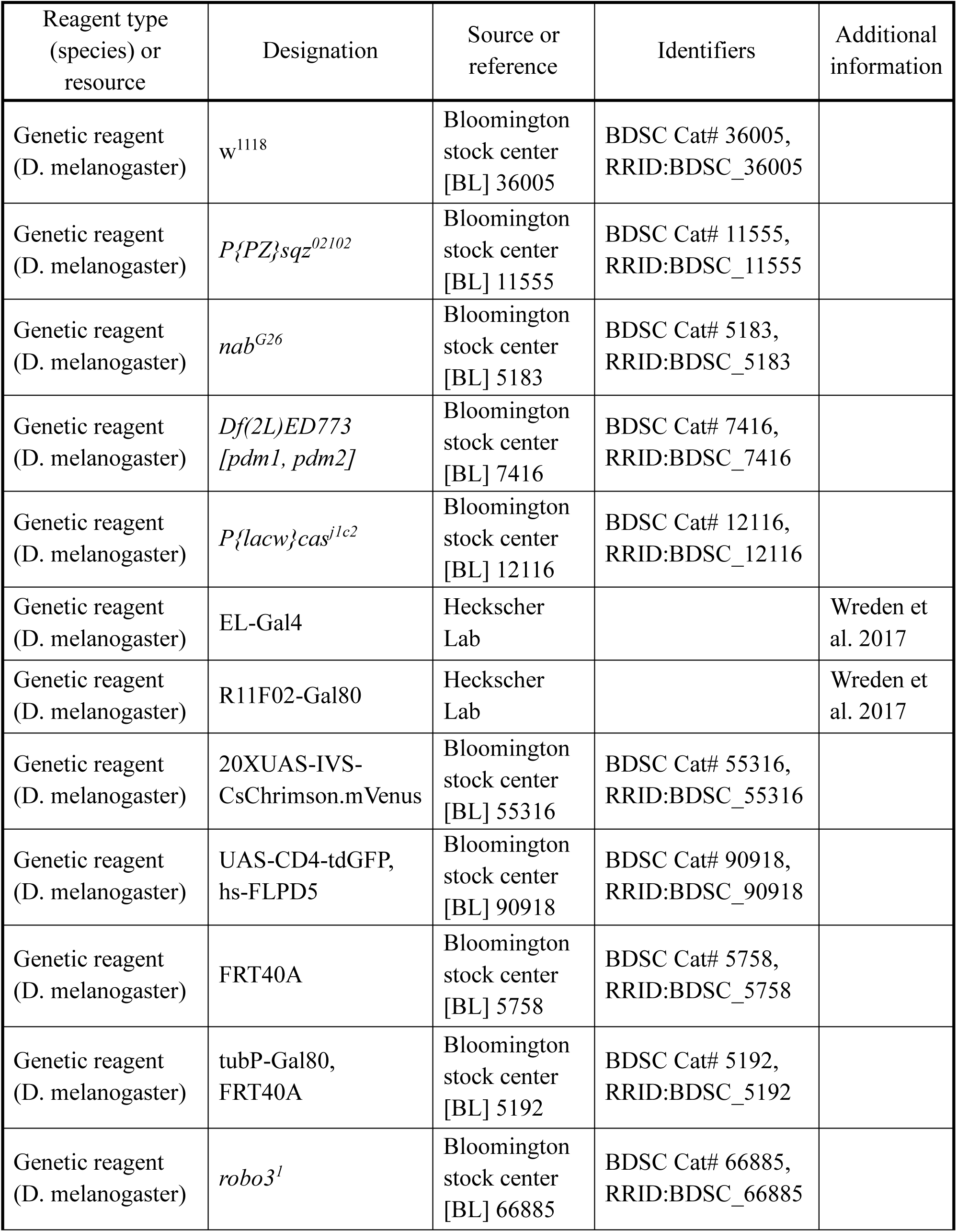
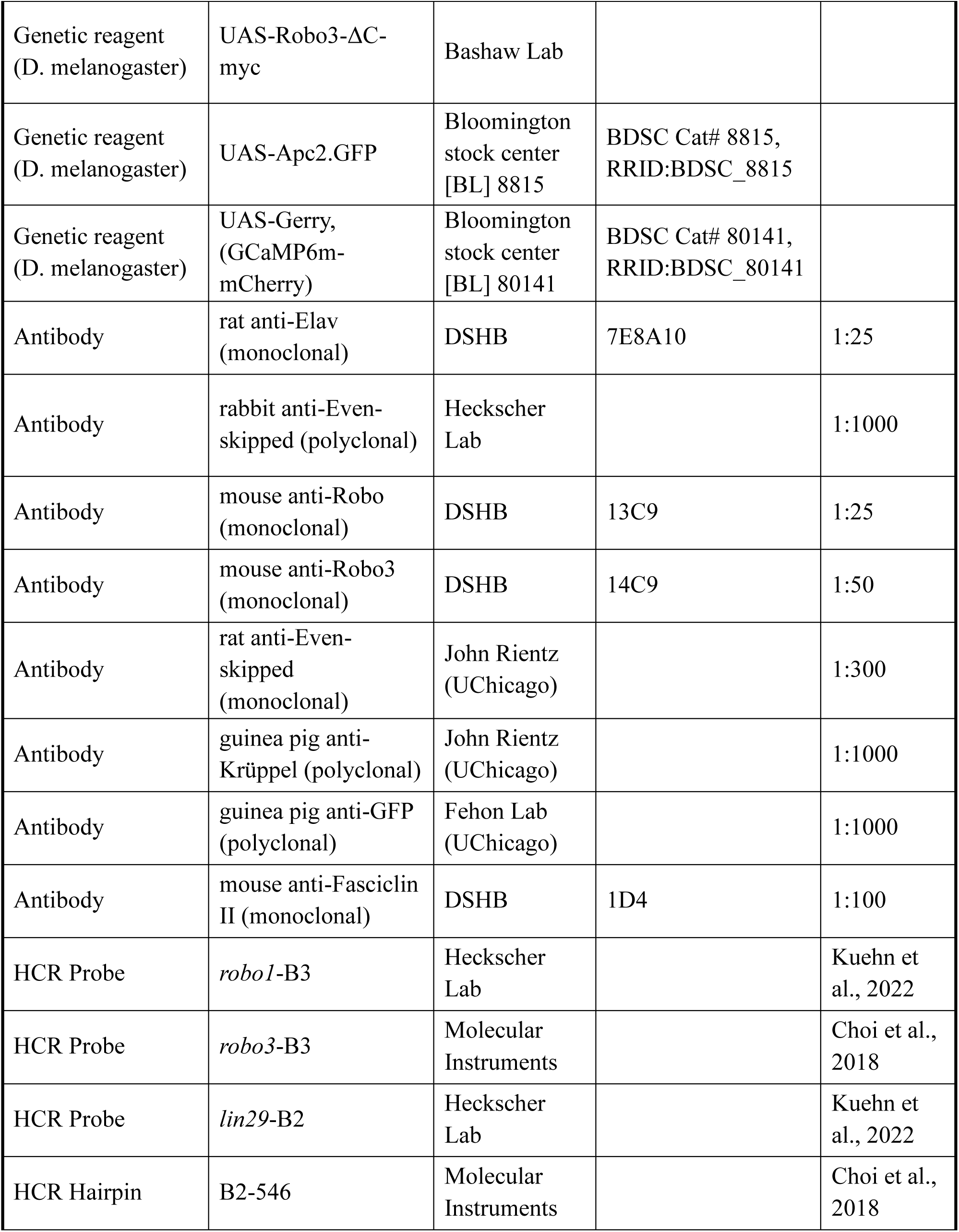
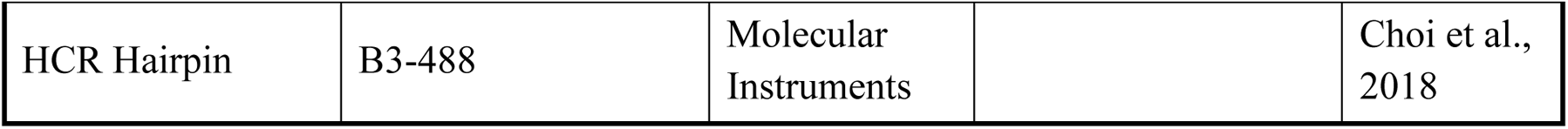

### Drosophila genetics

Standard methods were used for propagating fly stocks. For all experiments, embryos and larvae were raised at 25°C, unless otherwise noted. The following lines were used: w1118 (Bloomington Stock Center [BL] 36005), EL-GAL4 (Heckscher Lab [HL], {w[+mC]=EL-GAL4} (III)), R11F02-GAL80 (HL, P{w[+mC]=R11F02-GAL80}attP40), 10XUAS-IVS-myr::GFP (BL 32197), neoFRT40A (BL 5758), UAS-CD4-tdGFP, hs-FLPD5 (BL 90918), *robo3*^1^ (BL 66885), UAS-Robo3DN-myc (Robo3 dominant negative (II), gift of G. Bashaw), UAS-Apc2.GFP (BL 8815), UAS-Gerry (BL 80141), *cas^j^*^1c2^ (BL 12116), *Df(2L)ED773 [pdm1, pdm2]* (BL 7416), *sqz*^02102^ (BL 11555), *nab^G^*^26^ (BL 5183).

### Tissue preparation

Three tissue preparations were used for this research. First, whole mount embryos were prepared for imaging of neurogenic stages 10 through 16, during which antibodies can still penetrate the cuticle. Second, first instar larvae (L1) CNSs were isolated by dissecting the CNS from other larval tissue so that antibodies reach the CNS. Third, intact L1 larvae, in which the larvae and their CNS are maintained whole, were prepared for live imaging to study the function of neurons in response to sensory stimulation. For all preparations standard methods were used for fixation in fresh 3.7% formaldehyde (Electron Microscopy Sciences, Morgantown, PA) [15686]. For combined hybridization chain reaction and immunostaining samples were fixed twice; once before the HCR began and once after the HCR was finished. For L1 CNS isolations, L1 larvae were dissected in Baines solution containing 2 mM Ca^2+^ and 4 mM Mg^2+^ and adhered to poly-D-lysine (R&D Systems, Minneapolis, MN) [3439-100-01] coated coverslips. For calcium imaging, intact L1 larvae were selected and placed on a thin 1% agarose pad and restrained with a glass coverslip.

### HCR probe generation

Oligonucleotide HCR probes were ordered through Molecular Instruments or designed in-lab with either a B2 or B3 initiator sequence using previously tested code and parameters^87,88^ [Maximum percentage of C/G nucleotide content: 45-55%, maximum polyA/polyT homopolymer length: 5, maximum polyG/polyC homopolymer length: 3]. For any given number of probes that the code generated only 20 pairs were selected ensuring even coverage of the whole transcript and mitigating off-target promiscuity. The oligonucleotide sequences of interest were scanned for areas of somewhat similar sequences against the whole Drosophila Genome using NIH BLAST to prevent potential nonspecific binding. Areas of high sequence similarity were not considered for probe selection. For multiple transcript variants, probes were designed against the largest variant that encompassed all others. HCR probes were ordered through IDT Technologies. HCR probes were initially diluted in DEPC-treated water to a concentration of 100 μM.

### Immunostaining

Tissue was washed in phosphate buffered saline with 0.1% Tween-20 (PBST) and blocked for 1 hour at room temperature in 2% Normal Donkey Serum diluted in PBST. Samples were then incubated with primary antibodies overnight at 4°C and then with fluorescent secondary antibodies overnight at 4°C. Primary and secondary antibodies were diluted to their final concentrations in PBST. The following primary antibodies were used: anti-Eve (rabbit, 1:1000, Ellie Heckscher), anti-Eve (rat, 1:300, John Reinitz, University of Chicago), 1D4 anti-Fasciclin II (mouse, 1:100, DSHB), anti-GFP (guinea pig, 1:1000, Rick Fehon, University of Chicago), anti-Krüppel (guinea pig, 1:1000, John Reinitz, University of Chicago), 13C9 anti-Robo (mouse, 1:50, DSHB), 14C9 anti-Robo3 (mouse, 1:50, DSHB). All secondary antibodies were ordered from Jackson ImmunoLabs and used at a 1:400 dilution. Following staining, embryos were taken through a glycerol series of initial 25% and 50% glycerol diluted in DI-water (v/v) ending in a solution of 90% glycerol in DI-Water (v/v) and 4% n-propyl gallate (w/w). Embryos were kept in this final glycerol solution at 4°C until imaging.

### Hybridization chain reaction staining

Embryonic Samples were taken through HCR using published protocols (Henderson et al. 2024, Choi et al., 2018 [Molecular Instruments])^56,87^. Tissue was incubated with HCR probes in hybridization buffer (30% Formamide, 5X SSC, 9mM citric acid (pH 6.0), 0.1% Tween-20, 5% Dextran Sulfate) overnight at 37°C, then fluorescent hairpins in amplification buffer (5X SSC, 0.1% Tween-20, 5% Dextran Sulfate) overnight at room temperature in the dark. HCR probes used in this paper include: Robo1-B3 (Molecular Instruments), Robo2-B3 (MI), Robo3-B3 (MI), Lin29-B2 (HL). Fluorescent HCR hairpins were from Molecular Instruments and prepared according to published protocols: B3-h1-488, B3-h2-488, B2-h1-546, and B2-h2-546 (Molecular Instruments, Los Angeles, CA) [B3.0]. Embryos were either taken through an immunostaining protocol (see Immunostaining) or whole mounted in 90% glycerol with 4% N-propyl gallate for imaging. Whole mount embryos were staged for imaging based on morphological criteria.

### Single neuron MARCM assay

Single EL neurons that were genetically null for Robo3 were generated using a Mosaic Analysis with a Repressible Cell Marker (MARCM) assay^67^. The null Robo3 allele, robo3^1^, was combined with an FRT40A sit to be able to induce recombination with FLP expression. A heat-shock inducible FLP with a membrane GFP reporter UAS-CD4-tdGFP line was used to control the timing of recombination during embryonic development. For L1 larval labeling, embryos aged from 4 to 10 hours exposed to a 37C heat shock for 15 minutes and then allowed to develop to L1 larval stage. CNS tissue was dissected out and stained with anti-Eve, anti-GFP, and anti-Fas2. ELs were identified by their morphology.

### Calcium live imaging

Intact L1 larvae with early-born ELs expressing GCaMP and mCherry were selected for imaging and placed within a semi-restrained slide preparation^71^. We co-expressed GCaMP to monitor neural signaling and a calcium-insensitive mCherry to detect movement artifacts (Supplemental Figure 7). A speaker was placed next to the confocal stage to stimulate the mechanosensory circuit with a sound stimulus. Fluorescent activity was recorded for a total of 90 seconds(s) with 30s intervals before, during, and after the pure tone 1000 Hz sound stimulus. Image collection began simultaneously with playing the sound stimulus file. Following imaging larvae were removed from the preparation and allowed to recover on agar pad to control for animal health.

### Image acquisition

For fixed tissue images, data were acquired using a Zeiss LSM 800 confocal microscope with 40x oil (NA 1.3) or 63x oil (NA 1.4) objectives. For calcium imaging, data were acquired using a Zeiss LSM 800 confocal microscope with 40x oil (NA 1.3) objective using a 488nm and 546nm lasers with the pinhole completely open. Images were processed in ImageJ (NIH) and assembled in Adobe Illustrator.

### Image analysis

#### Robo1 and 3 transcript expression in ELs and U motor neurons

Late-stage embryos were stained with anti-Eve or anti-Elav, *robo1* or *robo3* HCR, and anti-Krüppel or *lin29* HCR and analyzed in ImageJ. The first abdominal segment was identified using tissue anatomy and Eve staining. Eve, Kr, and *lin29* were used to identify neuronal cell bodies, create regions of interest in ImageJ, and perform quantitative analysis of *robo1* and *3* expression using those ROIs. Because the cortex of the ventral nerve cord is a dense tissue, images were prepared in two ways. To highlight the general expression patterns of *robo1* and *robo3*, z-projections of hemisegments or cells of interest were created using Elav, Eve, Kr or *lin29* HCR to define the z-boundaries of the projection for Figures 1, 2, S1, and S2. To highlight the co-expression of *robo1* or *robo3* with Kr and *lin29*, Eve signal was used to create segmented z-projections with the image calculator tool in ImageJ for Figures 3, 4, and S3-S5.

#### Evaluation of early-born EL morphology

We used *EL-GAL4, R11F02-GAL80* to label early-born EL membranes with *UAS-mVenus*, express with or without *UAS-Robo3-ΔC-myc*, and evaluate the morphology in dissected L1 ventral nerve cords. L1 VNCs were co-stained with GFP, Fas2, and Eve antibodies and DAPI to confirm specificity of GFP labeling. Fas2 and DAPI staining were used to confirm position of neural morphology within the neuropil of the VNC. Fluorescence intensity of early-born EL membranes were quantified by performing line scans in ImageJ. Position of Fas2 and GFP signal were normalized across tissue by setting the bounds of each hemisegment neuropil with Fas2 signal. Fluorescence intensity of GFP signal was normalized within each tissue by calculating against background GFP signal. Plots of normalized spatial position versus fluorescence intensity were created using R Studio.

#### Evaluation of early-born EL dendrite arborization with Apc2.GFP

We used *EL-GAL4, R11F02-GAL80* to label early-born EL dendrites with *UAS-Apc2.GFP*, express with or without *UAS-Robo3-ΔC-myc*, and evaluate early-born EL dendrite arborization in dissected L1 ventral nerve cords. L1 VNCs were co-stained with GFP, Fas2, and Eve antibodies and DAPI to confirm specificity of GFP labeling and the boundaries of segment A1. Fas2 and DAPI staining were used to confirm position of dendritic signal within the neuropil of the VNC. With the neuropil identified, a ROI was created to evaluate each hemisegment. The number of GFP puncta per hemisegment were counted by thresholding, creating a binarized image of the GFP signal, and using the Analyze Particles function in ImageJ.

#### Live calcium imaging of early-born ELs

We used *EL-GAL4, R11F02-GAL80* to express *UAS-GCaMP6m.mCherry* with or without *UAS-Robo3-ΔC-myc* in intact L1 larvae. Following imaging the files were processed in ImageJ. ROIs were drawn around the GCaMP and mCherry signal within the A1 neuropil. For each frame of the recording fluorescence intensity was recorded for both the GCaMP and mCherry signal and then the fold change in GCaMP fluorescence intensity over time was normalized to the fold change in mCherry fluorescence intensity over time. Plots of the average normalized fold change and standard deviation were created from the data using R Studio. We also compared maximum change in fluorescence intensity between control and *UAS-Robo3-ΔC-myc* expressing larvae.

### Statistics

Descriptive statistics: average and standard deviation are reported. Each data point is plotted in every figure. Test statistics: All data sets were evaluated to determine if they followed a Gaussian distribution. In numerical data with two populations, we used either Welch’s t-Test for populations with unequal standard deviations or Mann-Whitney tests for populations with non-Gaussian distributions. In numerical data with three populations, we used Kruskal-Wallis test with Dunn’s multiple comparisons because all data sets had non-Gaussian distributions. Analysis completed using R Studio and GraphPad Prism 10.

## Acknowledgements

Rowan J Mangle, Sam Swank, Chris Wreden, Julia Meng, Marie Greaney, Deeptha Vasudevan, Ellen Lesser, Zarion Marshall, Rio Salazar, Annika Sharma, Grace Hu, Sarah Gagnon, Dr. Greg Bashaw, Dr. Robert Carrillo, Dr. Heather Marlow, Dr. Richard Fehon, FlyBase, Bloomington Drosophila Stock Center, Developmental Studies Hybridoma Bank, DBTG T32 HD055164 and NIH NINDS 1F31NS1S6008-01 to JEH, NIH R01-NS105748, NIH R56-NS134862, and MGCB start-up funds to ESH.

