## Supplemental Figures 1-5 for "Neural birth time and somatosensory circuit assembly are linked by Robo3 regulation of dendrite morphology"

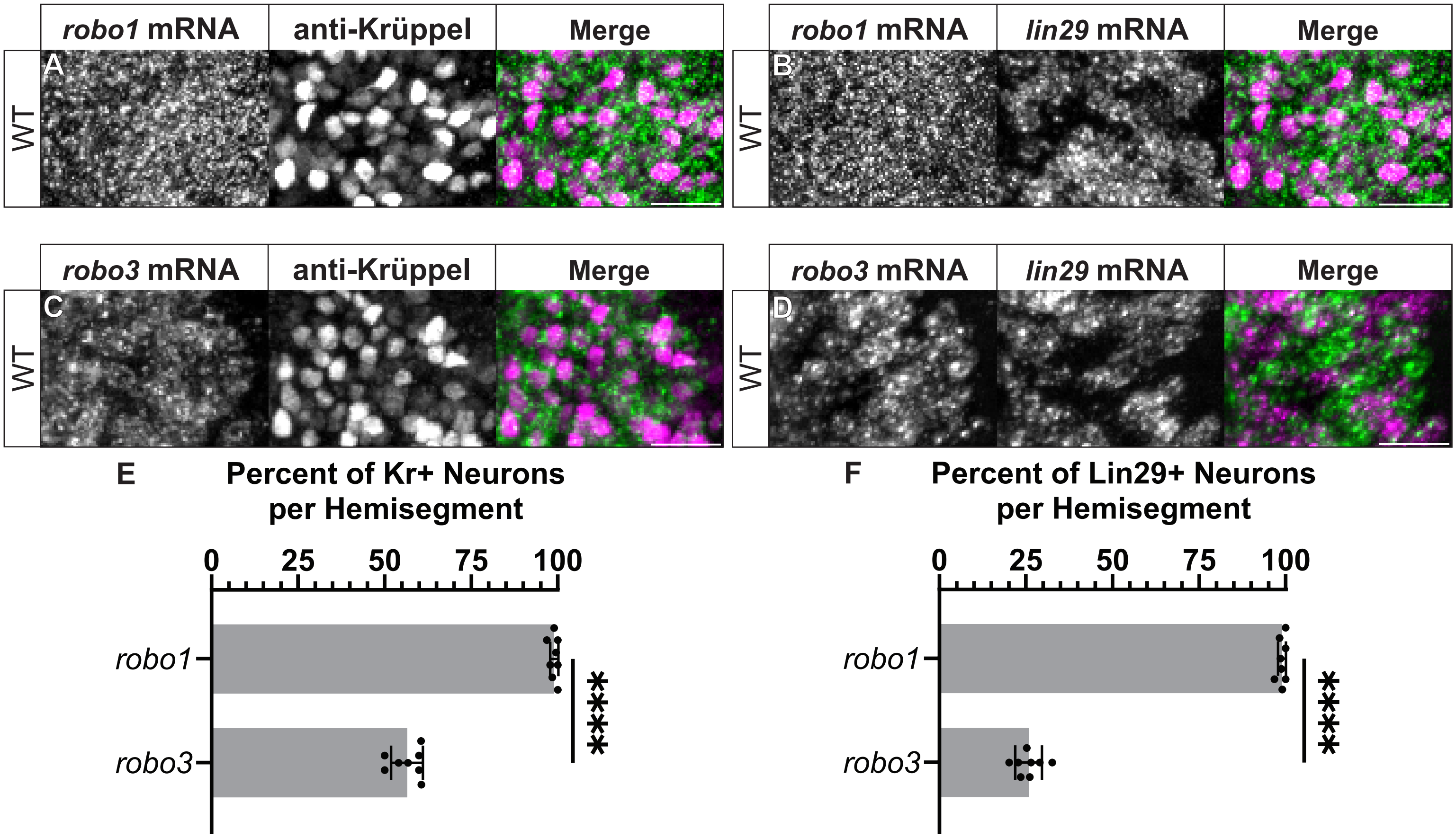

robo1 mRNA
anti-Krüppel
Merge
A
WT
robo1 mRNA
lin29 mRNA
Merge
B
WT
robo3 mRNA
anti-Krüppel
Merge
C
WT
robo3 mRNA
lin29 mRNA
Merge
D
WT
E
Percent of Kr+ Neurons per Hemisegment
0
25
50
75
100
robo1
robo3
****
F
Percent of Lin29+ Neurons per Hemisegment
0
25
50
75
100
robo1
robo3
****

**Supplemental Figure 1. Robo3 is found in a higher proportion of early-born vs late-born neurons.**

**A-D** Images of *robo1* and *robo3* mRNA expression compared to early-born marker (Kr) or late-born marker (Lin29). Single abdominal hemisegment is shown with anterior up and midline left. Scale bars represent 10 microns. **E-F** Quantification of Kr[+] or Lin29[+] neurons that co-express *robo1* or *robo3* mRNA. Each dot represents one hemisegment. Gray box shows average and black bars show standard deviation. Significance determined using an unpaired Welch's t-test, **** = $p<0.0001$.

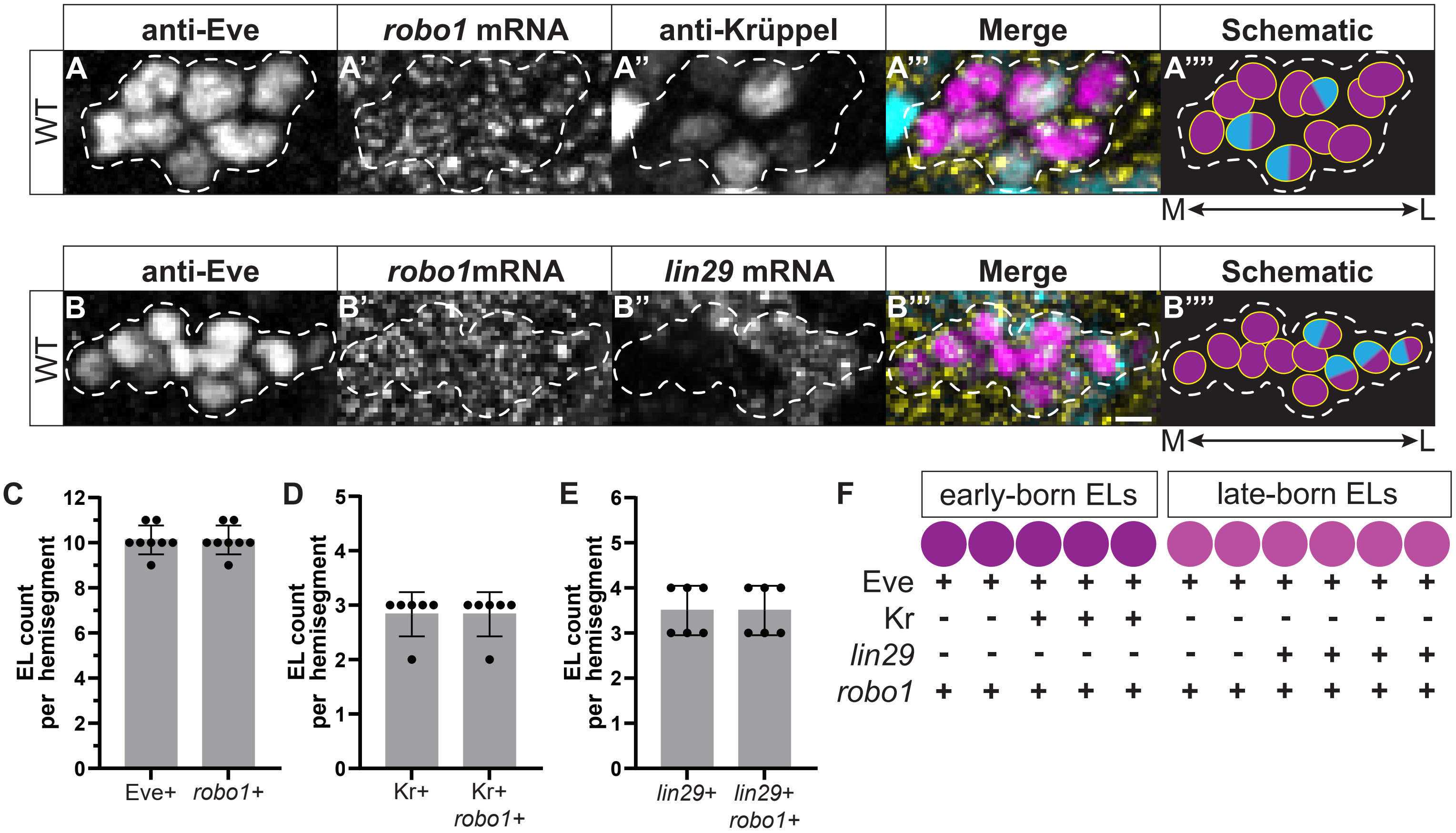

anti-Eve
robo1 mRNA
anti-Krüppel
Merge
Schematic
WT
A
A'
A''
A'''
A''''
M
L
anti-Eve
robo1mRNA
lin29 mRNA
Merge
Schematic
WT
B
B'
B''
B'''
B''''
M
L
C
EL count per hemisegment
Eve+
robo1+
D
EL count per hemisegment
Kr+
Kr+ robo1+
E
EL count per hemisegment
lin29+
lin29+ robo1+
F
early-born ELs
late-born ELs
Eve + + + + + + + + + + +
Kr - - + + + - - - - - -
lin29 - - - - - - - + + + +
robo1 + + + + + + + + + + +

**Supplemental Figure 2. Robo1 mRNA is expressed in both early-born and late-born ELs**

**A-B** Images and schematic summaries of ELs and *robo1* mRNA. The EL cluster is outlined by dashes. Images show a single abdominal hemisegment from a late-stage embryo, with anterior up and midline to the left. Scale bars represent 2.5 microns. In schematics, each circle represents one cell, with Eve in magenta, mRNA in yellow, and temporal marker in blue. **C-E** Quantifications of *robo1* mRNA expression patterns. Each dot represents measurement in one hemisegment. Gray box shows average and black bars show standard deviation. **F** Illustration of *robo1* mRNA expression in the NB3-3 lineage. Each circle is one cell: - represents no expression, + represents expression.

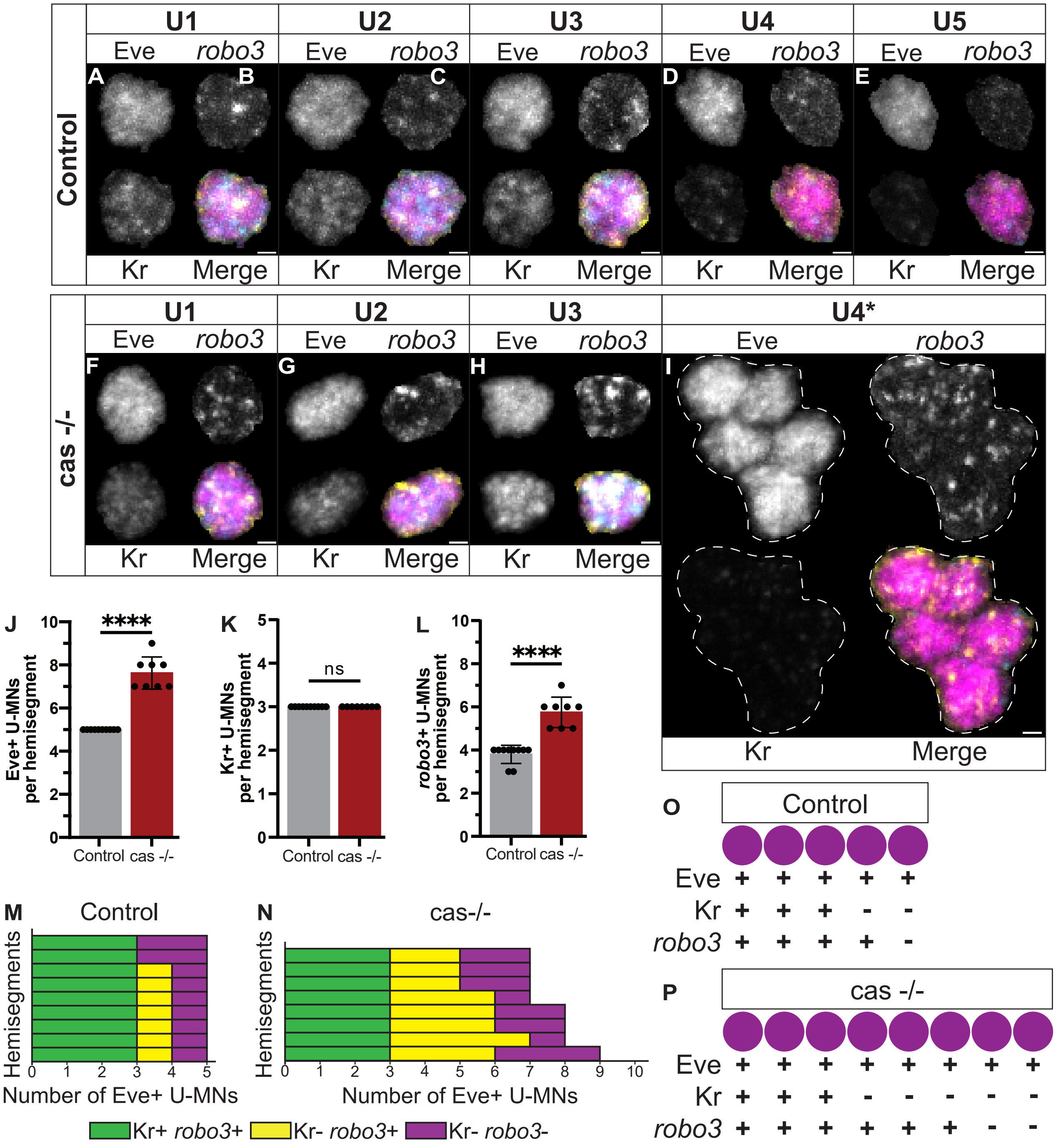
Control
U1
U2
U3
U4
U5
Eve
robo3
Kr
Merge
A
B
C
D
E
cas -/-
U4*
F
G
H
I
J
Eve+ U-MNs per hemisegment
****
Control cas -/-
K
Kr+ U-MNs per hemisegment
ns
L
robo3+ U-MNs per hemisegment
M
Control
Hemisegments
Number of Eve+ U-MNs
N
cas-/-
Kr+ robo3+
Kr- robo3+
Kr- robo3-
O
P
Eve
Kr
robo3
cas -/-

**Supplemental Figure 3. Robo3 is differentially expressed in U motor neurons associated with birth order.**

**A-I** Images of individual U-MNs, temporal identity, and their *robo3* mRNA expression. Representative cells in the abdomen of late-stage embryos are shown, with scale bars representing 1 micron. In K, the dashed line surrounds a cluster of Kr[-] U4-like motor neurons in cas homozygous mutant. **J-L** Quantification of expression patterns. Each dot represents the measurement in one hemisegment. Box shows average and black bars show standard deviation. Significance determined using Mann-Whitney test, ns = not significant, **** = $p<0.0001$. **M, N** Quantification of Kr and *robo3* co-expressoin in U-MNs. Each row represents one hemisegment. **O, P** Illustration of *robo3* mRNA expression in NB7-1 lineage in wild-type and *cas* mutant. Each circle is one cell: - represents no expression, an + expression.

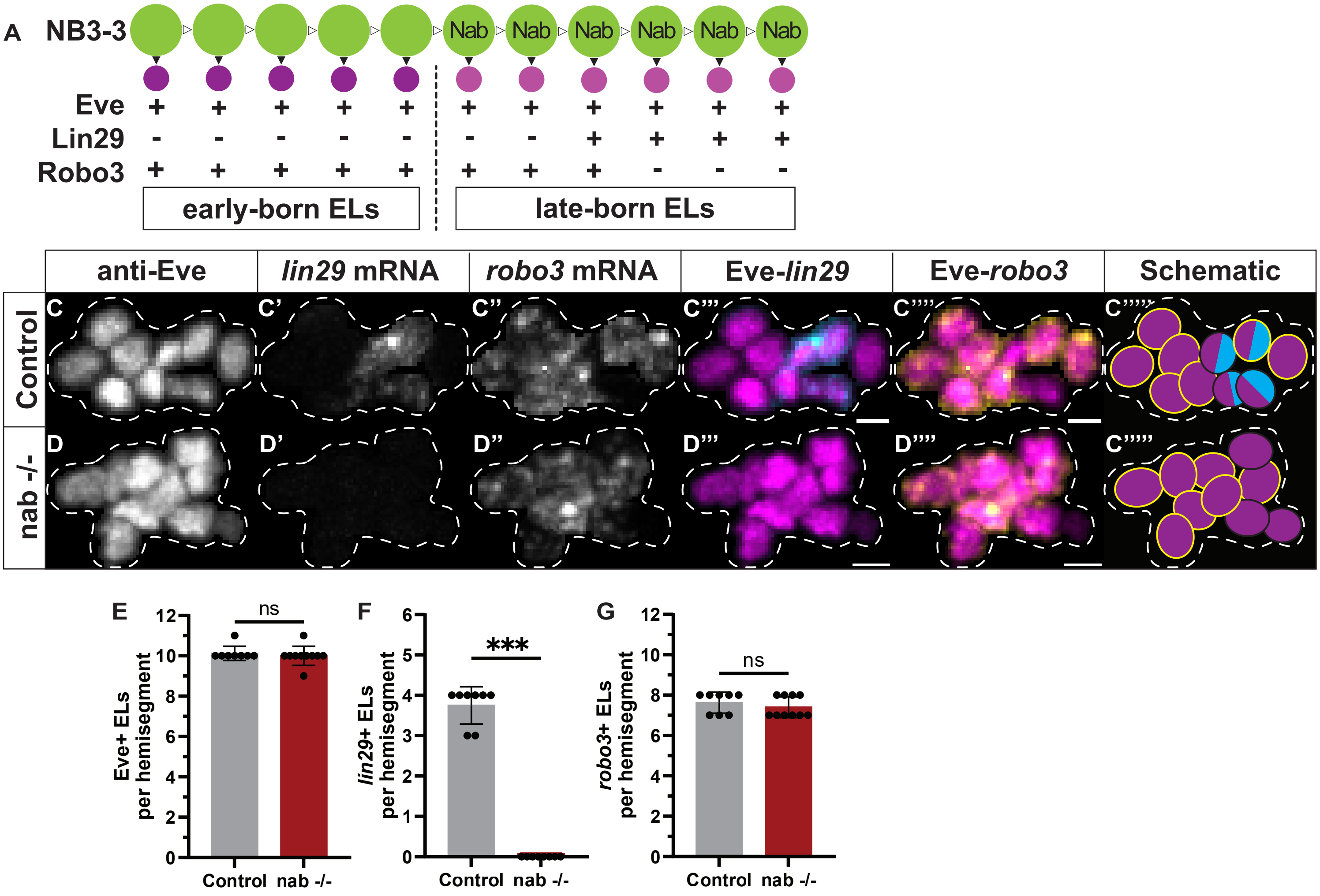
A
NB3-3
Nab
Eve
Lin29
Robo3
early-born ELs
late-born ELs
anti-Eve
lin29 mRNA
robo3 mRNA
Eve-lin29
Eve-robo3
Schematic
Control
nab -/-
C
C'
C''
C'''
C''''
C'''''
D
D'
D''
D'''
D''''
E
F
G
Eve+ ELs per hemisegment
lin29+ ELs per hemisegment
robo3+ ELs per hemisegment
ns
***
Control
nab -/-

**Supplemental Figure 4. Nab and Sqz regulate late temporal identity but not *robo3* expression in ELs**

**A-B** Images and schematic summaries of ELs, temporal identity, and *robo3* mRNA expression in ELs. EL cluster is outlined by dashes. Images who a single abdominal hemisegment from a late-stage embryo, with the anterior up and the midline left. Scale bars represent 2.5 microns. In schematics, each circle represents one cell, with Eve in magenta, the *robo3* mRNA in yellow, and Lin29 temporal marker in cyan. **C-E** Quantifications of expression patterns. Each dot represents the measurement in one hemisegment. Gray box shows average, and black bars show standard deviation. Significance determined using Mann-Whitney tests; ns = not signification, *** = p<0.001.

Genotypes: control is $cas^{j1c2}$ heterozygous, cas -/- is $cas^{j1c2}$ homozygous.

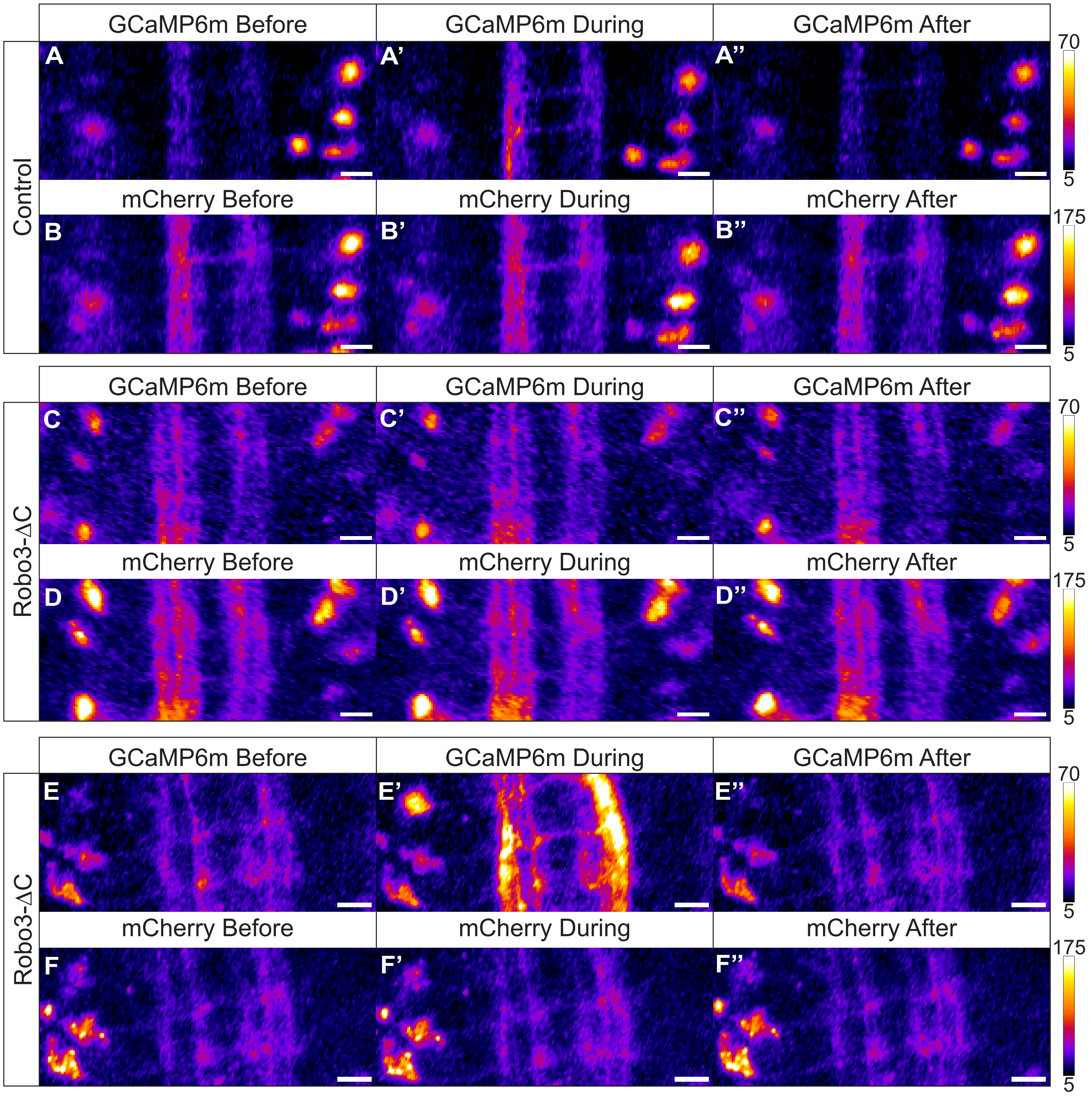
Control
Robo3-ΔC
Robo3-ΔC
GCaMP6m Before
GCaMP6m During
GCaMP6m After
mCherry Before
mCherry During
mCherry After
A
A'
A''
B
B'
B''
C
C'
C''
D
D'
D''
E
E'
E''
F
F'
F''
70
5
175
5

**Supplemental Figure 5. Robo3 loss of function does not affect Early-born EL expression of calcium insensitive mCherry.**

**A-F** Images from control and Robo3-ΔC early-born ELs expressing GCaMP6m:mCherry with one frame before, during and after stimulus. All images are from intact L1 larvae with anterior up from segment A1 to A3. Scale bars represent 10 microns
